# *Ex vivo* DHA supplementation suppresses prostaglandin E2 formation in primary human macrophages

**DOI:** 10.1101/2024.09.17.613409

**Authors:** Rebecca Kirchhoff, Nadja Kampschulte, Carina Rothweiler, Nadine Rohwer, Karsten-Henrich Weylandt, Nils Helge Schebb

## Abstract

**Scope:** There is evidence that intake of long-chain n-3 polyunsaturated fatty acids (PUFA) is associated with improved prognosis for inflammatory diseases. However, the underlying mechanisms are still subject of ongoing research. For this purpose, we developed an *ex vivo* n-3 PUFA supplementation strategy to test n-3 PUFA supplementation under controlled conditions in primary human macrophages.

**Methods and results:** Cells were supplemented with docosahexaenoic acid (DHA). Quality parameters to account for possible confounders were established for a reproducible and reliable supplementation. Following supplementation, PUFA pattern of cells was shifted towards a pattern reflecting that of subjects with a high n-3 PUFA status. This was accompanied by a decrease of arachidonic acid-derived oxylipins in a dose- and time-dependent manner in favor of n-3 PUFA ones. Stimulation with LPS resulted in decreased levels of pro-inflammatory prostaglandins in the DHA-supplemented cells, but no changes in cytokines.

**Conclusion:** *In vitro* supplementation studies with n-3 PUFA need rigorous controls to exclude background formation of oxylipins. By accounting for these possible confounders the desribed *ex vivo* approach is a promising tool for the mechanistic investigation of n-3 PUFA in primary human immune cells, offering an alternative for intervention studies in humans.

## Introduction

Dietary intake of n-3 and n-6 polyunsaturated fatty acids (PUFA) is essential for humans. In Western countries, n-6 PUFA are consumed in large amounts (e.g., by intake of plant oils from soy, sunflower, and corn oil) whereas the n-3 PUFA intake is low.^[1, 2]^ In addition, the conversion of alpha-linolenic acid to long-chain n-3 PUFA eicosapentaenoic acid (EPA) and docosahexaenoic acid (DHA) is low.^[3, 4]^ This is particularly the case because the n-6 PUFA linoleic acid competes for the same enzymes for its conversion to arachidonic acid (ARA).^[5]^ Thus, blood and tissue levels of EPA and DHA are low in humans on a Western diet,^[6, 7]^ indicated by a high n-6:n-3 PUFA ratio and high %n-6 in highly unsaturated fatty acid (HUFA) values, respectively.^[7, 8]^ These values are biomarkers for increased health risks, such as cardiometabolic events.^[9]^ A large number of studies shows that a high n-3 PUFA status, e.g. by consuming n-3 PUFA supplements, influences immune and inflammatory functions.^[10–12]^ However, the underlying mechanisms of action by which n-3 PUFA exert their anti-inflammatory effects are not fully understood.

Being key components of cell membranes, fatty acid (FA) composition impacts the fluidity of membranes which in turn influences the function of membrane-associated proteins.^[13]^ Moreover, PUFA undergo enzymatic and non-enzymatic oxidation giving rise to multiple oxylipins: Conversion by cyclooxygenases (COX) leads to highly potent prostaglandins (PG) and thromboxanes (Tx).^[14]^ Lipoxygenases (LOX) form regio- and stereoselective hydroperoxy-PUFA which are further reduced to hydroxy-fatty acids in the cells.^[15]^ Cytochrome P450 monooxygenases (CYP) form hydroxy-FA carrying the hydroxy-functionality at the ω and ω-n position or epoxy-FA. The latter can be easily hydrolyzed by the soluble epoxide hydrolase to *vic*-dihydroxy-FA. Non-enzymatic conversion by autoxidation leads to hydroperoxy-FA, hydroxy-FA and PG-like isoprostanoids.^[16]^

Several oxylipins are potent bioactive lipid mediators involved in the regulation of physiological processes such as fever and inflammation.^[17]^ ARA-derived PG formed by COX activity and leukotrienes formed by 5-LOX activity are well-investigated pro-inflammatory mediators.^[17]^ n3-PUFA reduce the conversion of n-6 PUFA and give rise to a distinct set of n3-PUFA-derived oxylipins. Several of those are discussed to have an anti-inflammatory effect, including 15-HEPE,^[18]^ or 7(*S*),17(*S*)-DiHDHA (resolvin (Rv)D5).^[19]^ Consequently, a shift in the oxylipin pattern is considered to be at least part of the anti-inflammatory mode of action of n-3 PUFA.

For elucidating the effects and the mode of action of n-3 PUFA, human intervention studies or clinical trials are ultimately neccessary.^[20–22]^ However, controlling nutrition and supplementation is challenging and thus, the results regarding anti-inflammatory effects on immune cells are inconsistent.^[23]^ Moreover, blood sampling and analysis of analytes such as oxylipins in clinical settings can lead to a large variations.^[24]^ In order to enable mechanistic investigation of immune cells without the need for human *in vivo* supplementation studies, we developed an *ex vivo* supplementation strategy for the detailed investigation of the physiological effects of n-3 PUFA and derived oxylipins in human macrophages. For all steps of the strategy, key parameters were identified and defined ensuring a reliable supplementation strategy. Applying the model to investigate the effect of the inflammatory stimulus LPS, changes in mRNA, protein and oxylipin levels of the supplemented cells were examined.

## Experimental section

### Chemicals and biological material

Human AB plasma was provided by the blood donation center at University Hospital Düsseldorf (Düsseldorf, Germany) and bought from BioIVT (West Sussex, United Kingdom) and Biowest (local distributor VWR, Darmstadt, Germany). Lymphocyte separation medium 1077 was obtained from PromoCell (Heidelberg, Germany). Recombinant human colony-stimulating factors CSF-1 (M-CSF), CSF-2 (GM-CSF), interferon (IFN)γ and interleukin 4 (IL-4) produced in *Escherichia coli* were purchased from PeproTech Germany (Hamburg, Germany). Ethylenediaminetetraacetic acid (EDTA) disodium dihydrate and dimethyl sulfoxide (DMSO) were obtained from Carl Roth (Karlsruhe, Germany). Docosahexaenoic acid (DHA) was bought from Nu-check Prep, Inc. (DHA > 99%) (Elysian, Minnesota, United States). Eicosapentaenoic acid (EPA) was obtained from Chemodex Ltd. (EPA ≥ 97%) (local distributor Biomol, Hamburg, Germany) and Nu-check Prep, Inc. (EPA > 99%). BCA reagent A was bought from Fisher Scientific (Schwerte, Germany). The ultra-pure water with a conductivity of >18 MΩ*cm was generated by the Barnstead Genpure Pro system from Thermo Fisher Scientific (Langenselbold, Germany). RPMI 1640 cell culture medium, L-glutamine and penicillin/streptomycin (5.000 units penicillin mL^-1^, 5 mg streptomycin mL^-1^), LPS from *E. coli* (0111:B4), dextran 500 from *Leuconostoc spp.* and copper sulfate pentahydrate were purchased at Merck (Schnelldorf, Germany).

### Cell cultivation

Primary human macrophages were prepared as described.^[25]^ In brief, peripheral blood monocytic cells (PBMC) were isolated from buffy coats obtained from blood donations at the University Hospital Düsseldorf. Blood samples were drawn with the informed consent of healthy human subjects. The study was approved by the Ethical Committee of the University of Wuppertal. PBMC were isolated from the supernatant after dextran (5%) sedimentation for 30–45 min. The erythrocytes in the remaining sediment were washed with PBS and subsequently frozen at −80 °C. Dextran supernatant was subjected to centrifugation (800 × *g* without deceleration, 10 min, 20 °C) on lymphocyte separation medium to isolate plasma and a leukocyte ring. Leukocytes were washed twice with PBS and seeded in 60.1 mm² dishes in RPMI medium (100 IU mL^-1^ penicillin and 100 µg mL^-1^ streptomycin (P/S), 2 mM L-glutamine) in a humidified incubator at 37 °C and 5% CO_2_ for 1 h. Non-adherent cells were collected by washing them off the dishes. Collected cells were washed two times with PBS and stored as dry pellets frozen at −80 °C until use. RPMI medium (P/S, 2 mM L-glutamine, and 5% human AB plasma) was added to the adherent cells. Monocytes were differentiated into macrophages using 10 ng mL^-1^ M-CSF for 7 days and additional 10 ng mL^-1^ IL-4 (M2-like) for the final 48 h.

### Supplementation experiments

Following pre-dilution in DMSO (50 mM), the n-3 PUFA preparation (DHA) was gently mixed with human plasma in order to stabilize the solution of PUFA: For medium supplemented medium, 20 µM n-3 PUFA preparation was added and for high supplemented medium 45 µM. Premixed plasma was added (5% (*v*/*v*)) to RPMI medium (P/S, 2 mM L-glutamine). Medium containing the same plasma, but without additional n-3 PUFA preparation, served as control medium with low n3-PUFA content (low). Macrophages were incubated with medium or high supplemented medium for either 2, 3 or 7 days. Every 2 to 3 days the cultivation medium was renewed. Prior to renewal of the medium, samples of supernatants were collected and frozen at −80 °C for analysis. For LPS stimulation, macrophages were treated with 1 µg mL^-1^ LPS 6 h before harvest. Macrophages were harvested by cold shock method,^[25]^ and cell pellets were frozen at −80 °C until analysis.

### Quantification of oxylipin, fatty acyl and protein levels by LC-MS/MS

Oxylipin determination was carried out in cell pellets, medium and human plasma as described.^[26–29]^ Briefly, cell pellets were resuspended in PBS, sonicated and protein content was determined via bicinchoninic acid assay.^[30]^ After addition of internal standards (IS), proteins were precipitated using MeOH (for non-esterified oxylipins) or *iso*-propanol (for total oxylipins) followed by centrifugation (20 000 × *g*, 10 min, 4 °C). For quantification of total oxylipin levels, supernatants were saponified using 0.6 M KOH in MeOH/H_2_O (75/25, *v*/*v*) for 30 min at 60 °C. Oxylipins were purified using solid phase extraction (SPE) and analyzed by means of LC–MS/MS.

The precipitated protein pellet following MeOH precipitation was analyzed for the abundance of key proteins.^[29, 31]^ Briefly, after resuspension in 5% (*w*/*v*) sodium deoxycholate and precipitation in ice-cold acetone, the pellet was redissolved in 6 M urea, disulfide bridges were reduced with dithiothreitol, and free sulfhydryl groups were alkylated with iodoacetamide. Following tryptic digestion for 15 h, and the addition of heavy labeled peptides, samples were subjected to SPE extraction and analyzed by LC-MS/MS.

The samples for oxylipin and peptide analysis were measured with separate methods on 1290 Infinity II LC systems (Agilent, Waldbronn, Germany), coupled to a QTRAP mass spectrometer (Sciex, Darmstadt, Germany) as described.^[26–29]^ Fatty acyl concentrations from the same samples as for the total oxylipins were determined by LC-MS/MS as described.^[32]^ Oxylipin, fatty acyl and peptide concentrations were quantified using external calibrations with IS and were calculated based on the total protein content in the cells. Fatty acyl levels in human plasma were analyzed by GC-FID following HCl-catalyzed transmethylation to fatty acid methyl esters (FAME).^[33]^

### RNA extraction and quantitative PCR analysis

Total RNA from M2-like macrophages was extracted using the RNeasy Mini Kit (Qiagen, Hilden, Germany) and QIAshredder spin columns (Qiagen) according to the manufacturer’s instructions. First-strand cDNA was synthesized with oligo (dT) primers and the iScript Select cDNA Synthesis Kit (Bio-Rad Laboratories, München, Germany). Gene expression was measured using the SsoFast EvaGreen Supermix on a CFX96 Real-Time PCR Detection System (both Bio-Rad Laboratories). Relative fold-changes of target gene expression were calculated by the comparative ΔCT method normalizing CT-values to the geometric mean of the CT-values of the housekeeping genes *RPL13A*, *SDHA* and *YWHAZ*. Primer sequences are listed in **Table S1**. qPCR measurements from the cell suspensions with and without sonication were comparable **(Figure S1)** allowing the parallel analysis of gene transcription, protein abundance and fatty acyl and protein levels.

## Results and Discussion

Dietary intake of n-3 PUFA is required for human health and sufficient intake of long-chain n-3 PUFA is associated with beneficial health effects.^[34, 35]^ In order to investigate the effects of these long-chain n-3 PUFA and their oxylipins, a reliable *ex vivo* supplementation strategy was developed allowing to change the cellular FA pattern of monocytes from subjects following a typical Western Diet to macrophages with a FA pattern comparable to subjects having a high n-3 PUFA status. This tool allows mechanistic investigation of the effects of long-chain n-3 PUFA in primary human immune cells under strictly controlled conditions.

### Characterization of key parameters for the ex vivo supplementation strategy

In order to enable a reproducible n-3 PUFA supplementation of primary human macrophages, several key paramters such as selection of buffy coats, plasma, n-3 PUFA preparation and composition of cell cultivation media have to be controlled **(Table 1)**.

**Table 1:**
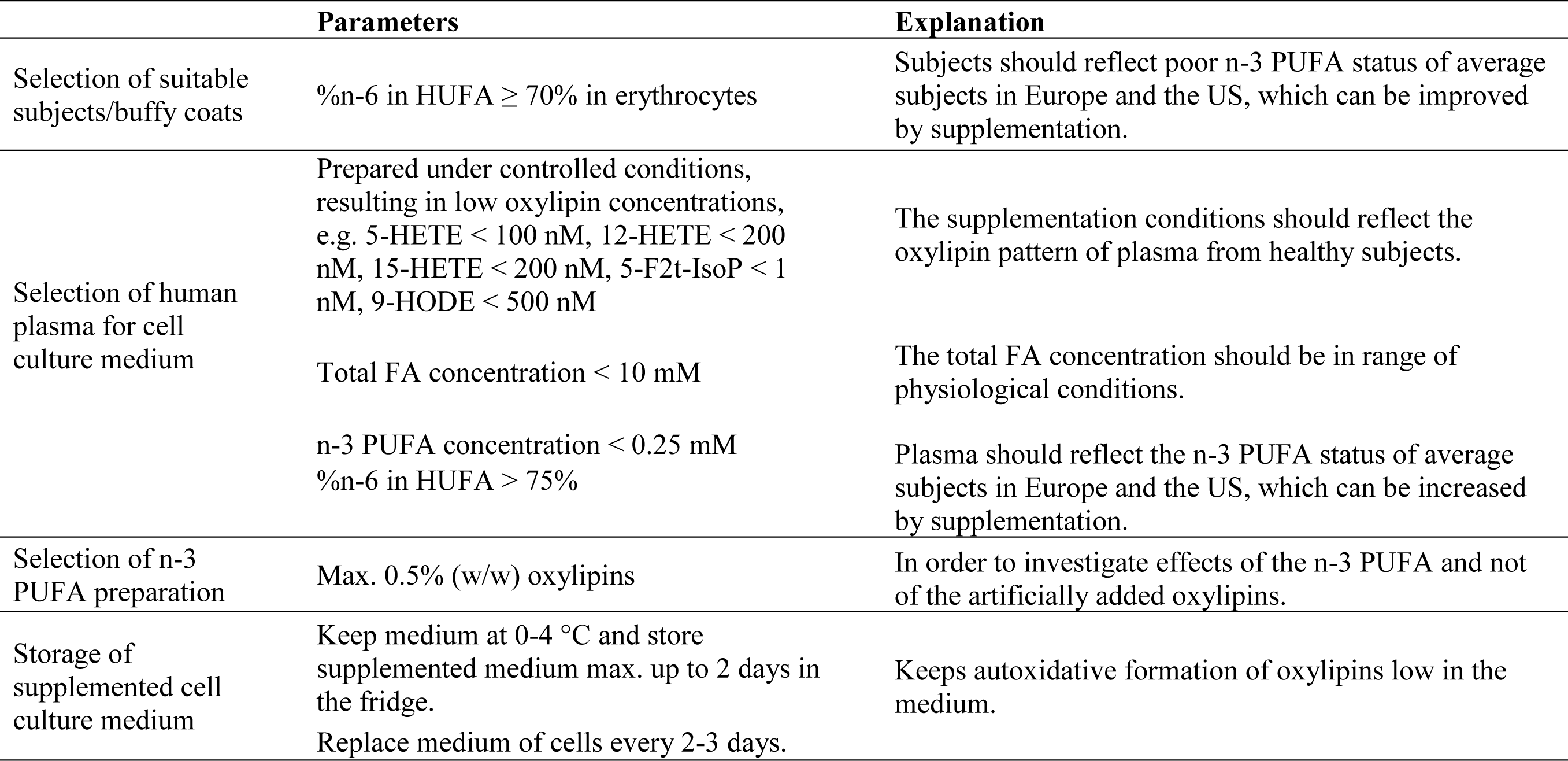
Key parameters for the selection of materials for the *ex vivo* supplementation of primary macrophages obtained from buffy coats.

|  | Parameters | Explanation |
| --- | --- | --- |
| Selection of suitable subjects/buffy coats | %n-6 in HUFA ≥ 70% in erythrocytes | Subjects should reflect poor n-3 PUFA status of average subjects in Europe and the US, which can be improved by supplementation. |
| Selection of human plasma for cell culture medium | Prepared under controlled conditions, resulting in low oxylipin concentrations, e.g. 5-HETE < 100 nM, 12-HETE < 200 nM, 15-HETE < 200 nM, 5-F2t-IsoP < 1 nM, 9-HODE < 500 nM | The supplementation conditions should reflect the oxylipin pattern of plasma from healthy subjects. |
|  | Total FA concentration < 10 mM | The total FA concentration should be in range of physiological conditions. |
|  | n-3 PUFA concentration < 0.25 mM<br>%n-6 in HUFA > 75% | Plasma should reflect the n-3 PUFA status of average subjects in Europe and the US, which can be increased by supplementation. |
| Selection of n-3 PUFA preparation | Max. 0.5% (w/w) oxylipins | In order to investigate effects of the n-3 PUFA and not of the artificially added oxylipins. |
| Storage of supplemented cell culture medium | Keep medium at 0-4 °C and store supplemented medium max. up to 2 days in the fridge. | Keeps autoxidative formation of oxylipins low in the medium. |
|  | Replace medium of cells every 2-3 days. |  |

#### a) Selection of cells from suitable subjects/buffy coats

FA status of the subjects (blood donors) was characterized based on the components of buffy coats. During the isolation of PBMC from buffy coats, fractions of erythrocytes, plasma and non-adherent cells were collected and investigated for their FA composition based on %n-6 in HUFA and n-6:n-3 ratio. Both markers of macrophages (e.g. subject 1: %n-6 in HUFA: 73.2 ± 2.4%; n-6:n-3 ratio: 3.2 ± 0.8) were reflected best by the erythrocytes (subject 1: %n-6 in HUFA: 72.8 ± 0.2%; n-6:n-3 ratio: 3.5 ± 0.1) **(Figure S2)**.

The PUFA pattern of erythrocytes, described as omega-3 index or %n-6 in HUFA, are established biomarkers for the individual n-3 PUFA status of subjects.^[9, 36, 37]^ They correlate with PUFA composition of tissues^[38, 39]^ as well as other blood cells.^[40]^ Due to the Western diet, typically containing high amounts of n-6 and low amounts n-3 PUFA,^[41]^ the average omega-3 index is low (4-6%) and %n-6 in HUFA is high (60-85%) in Europe and the US.^[9, 37,42]^ In order to reflect this, we only included subjects with > 70% n-6 in HUFA ensuring that they were representative of persons having a low n-3 PUFA status. Supporting the poor n-3 PUFA status of the German population, erythrocytes obtained from all buffy coats analyzed fulfilled the criterion (%n-6 in HUFA 72 ± 1 to 82 ± 1%) **(Table S2)**.

#### b) Selection of human plasma for cell cultivation media

Human plasma was selected based on the oxylipin and FA levels. A non-fasted plasma pool obtained from a local blood donation center and two commercially available plasma pools were compared for their oxylipin concentrations. Overall, oxylipin concentrations were up to 10-fold higher for the first and up to 600-fold higher in the second commercial plasma pools reaching the µM range for several oxylipins such as 5-HEPE or 14-HDHA **(Table S3)**. This can only be explained by inappropriate sampling and storage conditions during plasma preparation which leads to the artificial formation of oxylipins by both, non-enzymatic and enzymatic reactions.^[24, 27, 43]^ In order to reflect plasma conditions of healthy subjects,^[27]^ a plasma low in oxylipin levels has to be used for the cell culture medium. We defined concentrations of the five following oxylipins as a criterion if the plasma can be used for further experiments: 5-HETE (formed by autoxidation, 5-LOX-activity) < 100 nM, 12-HETE (formed during coagulation, 12-LOX activity) < 200 nM, 15-HETE < 200 nM (formed by autoxidation, 15-LOX activity), 5-F2t-IsoP < 1 nM (formed by autoxidation), 9-HODE < 500 nM (formed by autoxidation, oxylipin with the highest concentration in plasma).^[14, 44, 45]^

Plasma samples **(Figure S3)** from several healthy donors generated under controlled conditions at a local blood donation center were analyzed. FA concentration and pattern differed markedly **(Figure S3, Table S4)**: Due to different fasting states of the subjects, total FA concentrations were only about half as high in plasma B (7.7 ± 0.1 mM) and plasma E (7.1 ± 0.4 mM) compared to plasma C (15 ± 1 mM). n-3 PUFA concentrations were almost three times higher in plasma C and D (C, 0.48 ± 0.05 mM; D, 0.45 ± 0.02 mM) compared to plasma B (0.17 ± 0.02 mM) and %n-6 in HUFA ranged from 67 ± 1 (plasma D) to 83 ± 1 (plasma B). The FA pattern, e.g., n-6/n-3 ratio, n-3 index and %n-6 in HUFA was also different among the tested materials **(Table S4)** consistent with the strong variations reported for plasma even within a subject.^[40,46]^

For the selection of a plasma, we set the following criteria: i) Total FA concentration < 10 mM, ii) n-3 PUFA concentrations < 0.25 mM and iii) %n-6 in HUFA > 75%. Applying these criteria ensures that, i.) the addition of n-3 PUFA to plasma leads to FA levels which are in range of physiological total FA levels – considering plasma triglyceride values of maximum 3.3 mM,^[47]^ as most of FA are bound to triglycerides in plasma – and ii.) FA composition in plasma is comparable to plasma from subjects having a low n-3 PUFA status, typical for Europeans and US-Americans.^[9]^ Plasma B and E fulfilled all criteria and plasma B was used for the following experiments.

#### c) Selection of n-3 PUFA preparation

A comparison of commercially available n-3 PUFA preparations regarding their oxidation status revealed that oxylipin content differed more than 10-fold **(Table S5, S6)**. Different extraction and purification methods, but particularly formulation, storage as well as packaging conditions influence autoxidation of the PUFA and thus, could explain different amounts of oxylipins. Considering that concentrations up to 45 µM of the n-3 PUFA preparation are used in the cell culture medium, this could lead to relevant oxylipin concentrations in the medium: For example, using a commercial n-3 PUFA preparation with 3% (*m*/*m*) oxylipins, 18-HEPE concentrations in the supplemented medium were more than 50-fold higher (59 nM) than in the non-supplemented medium. This makes it impossible to differentiate whether biological effects originate from the n-3 PUFA or the artificially added oxylipins. Thus, only using a n-3 PUFA preparation with < 0.5% oxylipins can ensure that mainly effects of the n-3 PUFA are investigated and not of the artificially formed and none-intentionally added oxylipins, as several of those are potent bioactive lipid mediators.

#### d) Composition and storage of n-3 PUFA-supplemented cell culture media

Addition of DHA led to approximately five- (medium, 20 µM DHA added) or ten-fold (high, 45 µM DHA added) increased DHA concentrations compared to the non-supplemented medium (low). Compared to ARA (22 ± 1 µM), the resulting DHA concentration was equal (medium, 21 ± 1 µM) or twice as high (high, 41 ± 2 µM) **(Figure 1A (i))**. %n-6 in HUFA was decreased from 81.0 ± 0.6% (low) to 53.9 ± 1.8% (medium) or 39.4 ± 0.4% (high) **(Figure 1A (ii))**. In line with this, n-6:n-3 ratio ≥ C18 was decreased from 17.4 ± 0.8 (low) to 5.5 ± 0.3 (medium) or 3.1 ± 0.1 (high) and to a similar extent for n-6:n-3 ratio > C18 in both supplemented media **(Figure 1A (ii))**.

**Figure 1:**
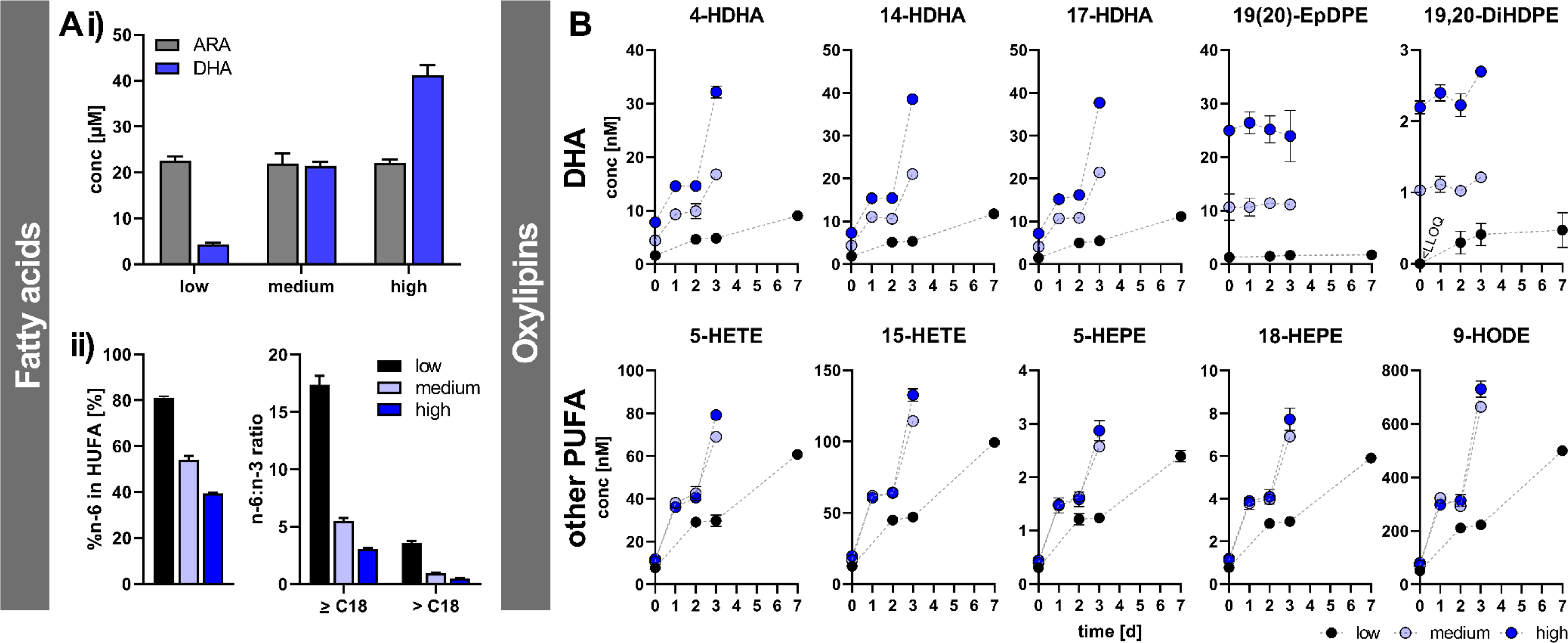
Characterization of n-3 PUFA-supplemented cell culture media. Media were prepared by using plasma B and addition of DHA preparation (> 99%) to the cell culture medium leading to concentrations of 4 µM (low, non-supplemented control), 21 µM (medium) or 41 µM (high) DHA in the cell culture media. **A) (i)** Total FA concentrations were analyzed in the different media by means of LC-MS/MS. **(ii)** %n-6 in HUFA was calculated from concentrations of C20:3 n-6, C20:4 n-6, C22:4 n-6, C22:5 n-6, C20:3 n-9, C20:5 n-3, C22:5 n-3, C22:6 n-3. n-6:n-3 ratios were determined by FA concentrations of C20:4 n-6, C20:5 n-3 and C22:6 n-3. n-6:n:3 ratio ≥ C18 includes additionally concentrations of C18:2 n-6 and C18:3 n-3. **B)** Total oxylipin concentrations were analyzed at baseline and after incubation for 1, 2, 3 and 7 days at 37 °C by means of LC-MS/MS. Results are shown as mean ± SD, *n* = 3.

The %n-6 in HUFA of non-supplemented medium is comparable to plasma %n-6 in HUFA from subjects having a diet low in n-3 and high in n-6 PUFA such as the Western Diet in the US and Europe.^[9]^ Thus, incubation of cells with a non-supplemented medium resembles the nutritional status of subjects low in n-3 and high in n-6 PUFA.

Medium supplemented medium with 54 ± 2% n-6 in HUFA is similar to human plasma after supplementation with DHA in human nutrition studies^[20, 48]^ and in populations whose diets are rich in fish.^[9]^ The FA composition in high supplementation medium with DHA concentrations twice as high as ARA resulting in 39 ± 0.4% n-6 in HUFA is slightly higher than what can be reasonably achieved by diet in plasma. However, %n-6 in HUFA values low as 30% have been reported in erythrocytes and human tissues, e.g., for subjects from Greenland.^[9, 49]^

As PUFA are prone to autoxidation, we investigated the stability of cultivation medium after preparation and after incubation under cell culture conditions or in the fridge. At baseline, all DHA-derived oxylipins were higher concentrated in the supplemented media compared to the non-supplemented (e.g. 4-HDHA: low, 1.62 ± 0.03 nM; medium, 4.4 ± 0.2 nM; high, 7.8 ± 0.3 nM), whereas oxylipins derived from other PUFA such as ARA or EPA were the same in all media (e.g. 5-HEPE: low, 0.31 ± 0.03 nM; medium, 0.40 ± 0.03 nM; high, 0.44 ± 0.04 nM) **(Fig. 1B)**. This indicates that the majority of DHA-derived oxylipins in supplemented media originated from the PUFA preparation, as plasma contributed only a small part of total DHA oxylipins **(Figure S4)** and oxylipin pattern of PUFA preparation matched those found in the media **(Table S6, Figure S4)**.

When medium was stored at 37 °C for several days, monohydroxy-FA concentrations increased over time, whereas total FA concentrations, epoxy-FA and *vic*-dihydroxy-FA were not affected **(Figure 1B)**. The slope of oxylipin formation was dose-dependently higher with increasing amounts of supplemented n3-PUFA in the medium, as the rate of autoxidation is mainly dependent on i) the number of double bounds being prone to abstraction of a bis-allylic hydrogen atom and ii) the number of hydroperoxides, which are contained or formed in the PUFA preparation **(Table S6)**, leading to radicals and starting the chain reaction.^[50]^ After three days of storage, oxylipin concentrations increased linearly by approximately 3-4-fold in the control medium and 6-9-fold in the supplemented media **(Figure 1B)**. In order to avoid this massive formation of monohydroxy-FA, supplemented medium was replaced after 48 h of cell cultivation. Of note, supplemented medium stored in the fridge at 4 °C underwent autoxidation resulting in oxylipin levels comparable to storage at 37 °C (e.g. for 14-HDHA in high supplemented medium stored for 4 days: 4 °C, 13.8 ± 0.4 nM; 37 °C, 15.4 ± 0.3 nM) **(Figure S5)**. For this reason, also supplemented media should be prepared freshly and stored at 0-4 °C as short as possible but at most up to two days.

Overall, several key parameters such as the selection of buffy coats, plasma and n-3 PUFA preparation have to be controlled for an *ex vivo* n-3 PUFA supplementation of primary macrophages **(Table 1)**. Only then it can be ensured that effects of the n-3 PUFA are investigated and not of the artificially formed and non-intentionally added oxylipins.

### Effects of n-3 PUFA supplementation on macrophages FA pattern

Following supplementation of macrophages for two, three or seven days with the DHA-supplemented medium, the FA profile as well as the oxylipin pattern were strikingly changed in the macrophages compared to non-supplemented cells (low) **(Figure 2)**: Non-supplemented cells showed 84 ± 2% n-6 in HUFA which is comparable to values found in tissues from subjects having a low n-3 PUFA status.^[9]^ In supplemented macrophages, %n-6 in HUFA was dose- and time-dependently lower (after two-day supplementation: medium, 51 ± 2%; high, 35 ± 2%) **(Figure 2A)**. The change in %n-6 in HUFA was strongest during the first two days of supplementation, then the decrease flattened with duration of supplementation indicating that the cells are approaching steady-state. After seven days of supplementation %n-6 in HUFA was decreased to a minimum of 23 ± 1% or 14 ± 1% in the cells incubated with medium or high supplemented medium, respectively.

**Figure 2:**
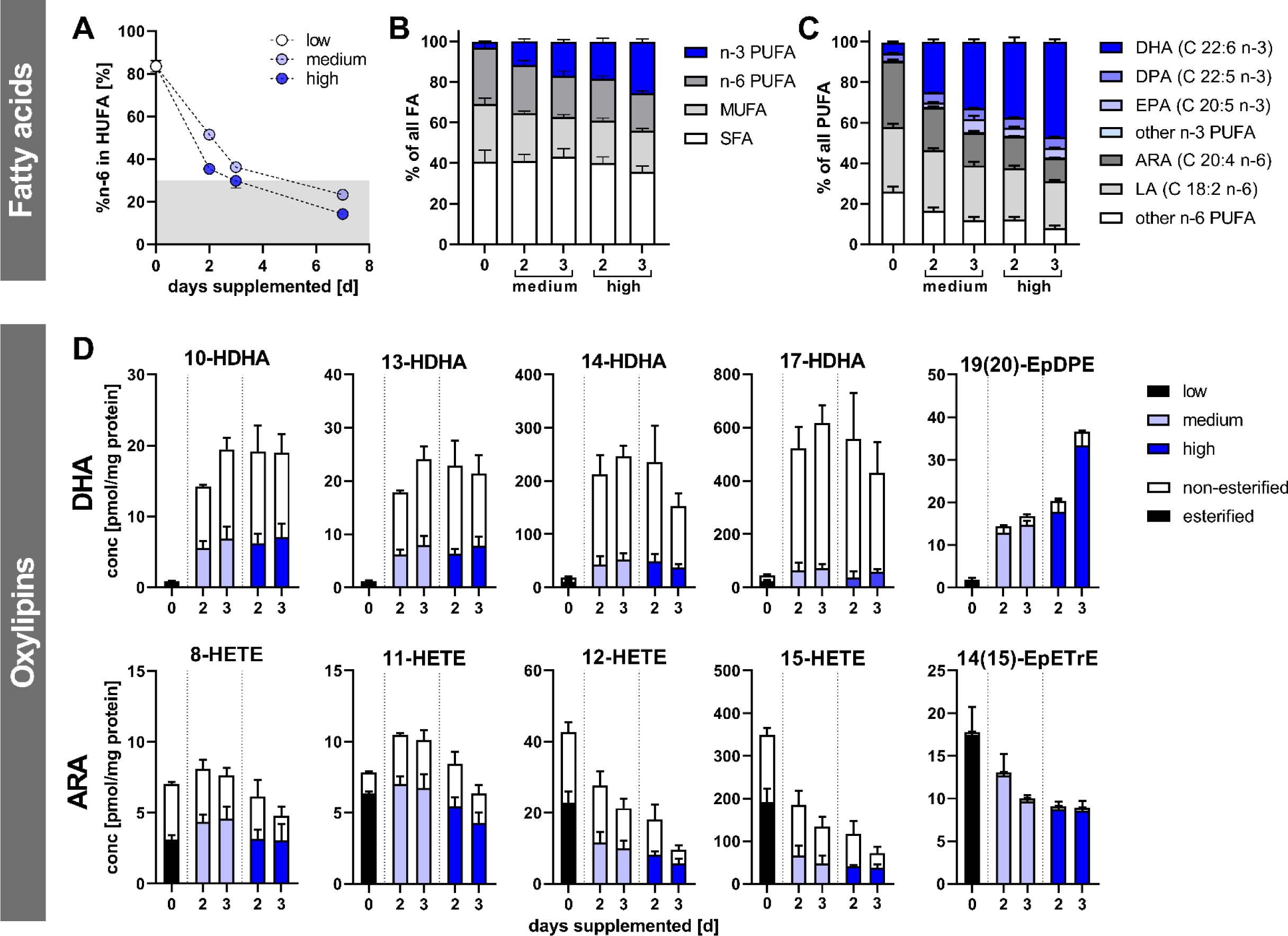
Characterization of FA and oxylipin pattern in macrophages after supplementation with n-3 PUFA. After isolation of monocytes and differentiation into M2-like macrophages, cells were supplemented using media containing 4 µM (low, non-supplemented control), 21 µM (medium) or 41 µM (high) DHA for 2, 3 or 7 days. Cell pellets were analyzed for total FA and non-esterified and total oxylipins by means of LC-MS/MS. **A)** %n-6 in HUFA was calculated from total FA concentrations of C20:3 n-6, C20:4 n-6, C22:4 n-6, C22:5 n-6, C20:3 n-9, C20:5 n-3, C22:5 n-3, C22:6 n-3. **B)** Relative distribution of n-3 PUFA, n-6 PUFA, MUFA and SFA. **C)** Relative distribution of PUFA including C22:6 n-3, C22:5 n-3, C20:5 n-3, C18:3 n-3 and C22:2 n-6, C22:4 n-6, C22:5 n-6, C20:3 n-6, C20:4 n-6, C18:2 n-6, C18:3 n-6. **D)** Concentrations of selected, most abundant oxylipins derived from DHA and ARA in the cells. Results are shown as mean ± SEM, *n* = 3-6.

Human nutrition intervention studies reduced %n-6 in HUFA to about 50-60% in erythrocytes after three to five months by daily doses between 1-2 g.^[20–22]^ These %n-6 in HUFA values achieved by oral supplementation are comparable with those of the macrophages supplemented with DHA in the cultivation medium (medium) for 2 days (51 ± 2%).

Taking a closer look at the relative FA pattern of the supplemented cells, the relative amount of saturated fatty acids (SFA), monounsaturated fatty acids (MUFA) and PUFA was unchanged independently from the n-3 PUFA status **(Figure 2B)**. Thus, supplementation leads to the replacement of n-6 PUFA by n-3 PUFA (increase of n-3 PUFA from 2.9 ± 0.7% to approximately 12-25% in a dose- and time-dependent manner). This is consistent with supplementation studies in humans and animals^[20–22, 38]^ and mainly driven by an increase in DHA (from 1.7 ± 0.3% to 8.9-21%) and a decrease in ARA (from 13-16% to 9.8-13% of all FA) **(Fig. 2 C)**. Though only DHA was supplemented, EPA was also higher in the supplemented cells (0.8-2.2% of all FA) compared to the control cells (0.09 ± 0.01% of all FA). An increase of EPA in response to DHA supplementation is discussed to be due to a formation of EPA by retroconversion from DHA^[20, 51, 52]^ or due to a slower EPA metabolism in the presence of high DHA levels.^[48]^ Recent studies show that DHA inhibits its own synthesis at the level of EPA:^[53]^ the higher EPA level could also result from a change in synthesis rates of EPA/DHA from α-linolenic acid. Even though the molecular mechanism is still a matter of debate, an increase of EPA in blood and tissue following DHA-only supplementation is commonly described in humans and animal models.^[20, 48, 52, 54, 55]^ The decrease of ARA is similar to previous reports for erythrocytes in human intervention studies, where ARA was decreased from approximately 13-16% to 9.8-13% of all FA following n-3 PUFA supplementation.^[20–22]^

In summary, we showed how the PUFA pattern of monocytes from subjects having a low n-3 PUFA status can be modified to macrophages resembling a PUFA pattern with a high n-3 PUFA status under controlled conditions. Moreover, higher n-3 PUFA levels can be achieved when supplementing at a higher dose, allowing the detailed investigation of effects of an increased n-3 PUFA status in human immune cells.

### Effects of n-3 PUFA supplementation on the oxylipin pattern of macrophages

The oxylipin pattern was changed in the supplemented macrophages compared to control cells (low) in a similar manner as their PUFA precursors **(Figure 2C-D)**: While DHA was increased up to 10-fold, DHA-derived oxylipins increased 10-20-fold, whereas ARA-derived oxylipins decreased about 2-fold in the supplemented cells. Our findings, (i) an increase in n-3 PUFA-derived oxylipins while ARA-derived ones decreased, (ii) the relative increase in n-3 PUFA concentrations was higher than the decrease in ARA concentrations, and (iii) that the relative change in oxylipins was higher than that of the corresponding PUFA by supplementation, are consistent with nutrition intervention studies.^[38, 56–58]^

Interestingly, concentrations of non-esterified monohydroxy-DHA were increased to a greater extent by supplementation (approximately 5-20 times more) than esterified oxylipins **(Figure 2D)**. In contrast, the ratio of unbound to esterified oxylipins was hardly changed for ARA-derived oxylipins in the supplemented cells. This could indicate that formation or uptake of DHA-derived oxylipins from the medium was more rapid than their incorporation into lipids such as phospholipids of the cell membrane.

Highest total oxylipin concentrations were found for 17-HDHA in the supplemented cells (430 ± 120 – 620 ± 81 pmol mg^-1^ protein), whereas 15-HETE was the most abundant oxylipin in the non-supplemented cells (low, 349 ± 73 pmol mg^-1^ protein) **(Figure 2D)**. 14- and 17-HDHA concentrations were increased strongest by supplementation (difference between two-day supplementation with medium supplemented medium and low: 14-HDHA, 194 ± 51 pmol mg^-1^ protein; 17-HDHA 475 ± 102 pmol mg^-1^ protein), whereas others such as 10-HDHA were less affected (13.3 ± 31.2 pmol mg^-1^ protein). 12- and 15-HETE were dose- and time-dependently lower in the supplemented cells, whereas other monohydroxy-ARA such as 8-HETE were not altered by supplementation **(Figure 2D)**.

The observed changes in oxylipin concentrations are likely due to the modified PUFA profile and reulting enzymatic conversion. To investigate this, we compared oxylipins in the medium following a two-day supplementation period with medium which was stored under identical conditions, but without cells reflecting autoxidation **(Figure 3A, Figure S6A)**: Monohydroxy-PUFA were lower in the cells’ supernatant than in the medium without cells under all conditions **(Figure 3A)**. The highest difference was observed for high supplementation medium (∼ 6-10 nM), whereas the difference was smaller in the non-supplemented medium (∼ 1-3 nM). However, - within the supplementation groups (low, medium, high) - concentration differences were comparable for all monohydroxy-FA from DHA (e.g., low: 7-HDHA, 1.0 nM, 17-HDHA, 1.0 nM; high: 7-HDHA, 5.9 nM, 17-HDHA, 6.6 nM). As the original concentrations of the various hydroxy-FA are of the same order of magnitude, **(Figure 3A, Tabel S6),** this suggests, that hydroxy-FA are unselectively taken up into the cells in a dose-dependent manner.

**Figure 3:**
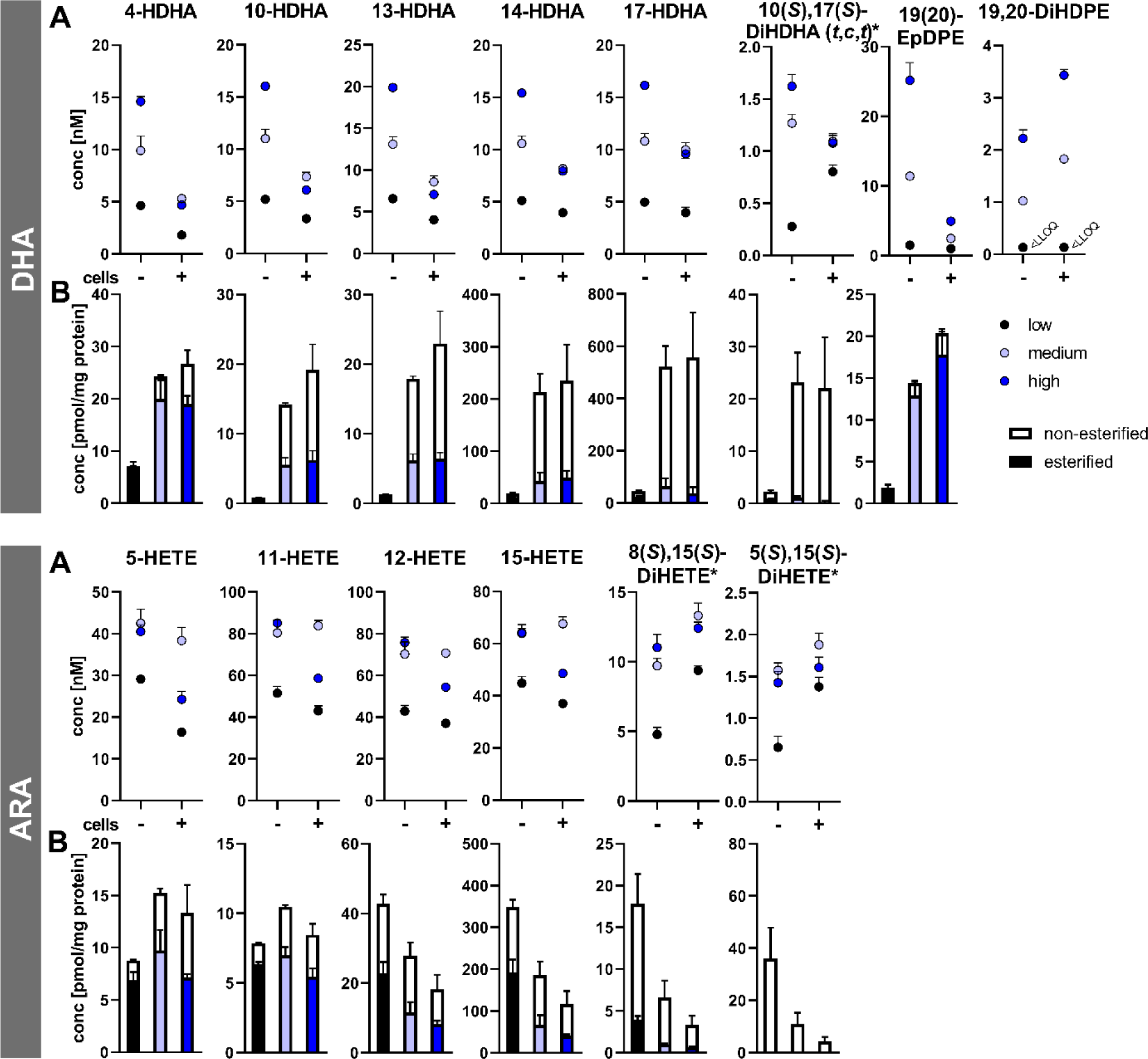
Oxylipin pattern of A) medium (without (−) and with cells (+)) and B) the macrophages after two days of incubation with different levels of DHA. Monocytes were isolated from buffy coats and differentiated into M2-like macrophages. Incubations were carried out using media containing 4 µM (low, non-supplemented control), 21 µM (medium) or 41 µM (high) DHA for 2 days with and without cells. Cell culture media was analyzed for total oxylipins, and cell pellets for total and non-esterified oxylipins by means of LC-MS/MS. Results are shown as mean ± SD (medium) or mean ± SEM (cells), *n* = 3. (*for **A)**, oxylipin and enantiomer)

In the cells, DHA-derived oxylipins were increased differently by supplementation, e.g. 17-HDHA was increased 25-fold stronger than 4-HDHA (18 ± 6 pmol mg^-1^ protein vs. 490 ± 200 pmol mg^-1^ protein) **(Figure 3B top)**. This could be explained by enzymatic conversion of DHA by 15-LOX-1 activity, which is highly abundant in M2-like macrophages.^[29]^ 15-LOX-1 has a dual reaction specificity which gives rise to both 17- and 14-HDHA.^[59, 60]^ This explains the high concentrations of the second most abundant oxylipin in the supplemented cells, 14-HDHA. Similar findings were observed for 15-LOX-1 products of EPA, 12- and 15-HEPE **(Figure S6)**. 12- and 15-HETE were both dose-dependently lower in supplemented cells **(Figure 3B bottom)** suggesting a lower conversion of n-6 PUFA by competition with supplemented DHA. This is supported by the finding, that 15-LOX-1 preferentially converts DHA and EPA compared to ARA.^[59]^ This also explains the relative higher increase in 14- and 17-HDHA compared to the elevation of DHA following supplementation.

The dual reaction specify of 15-LOX-1 results in n-6 and n-9 products (i.e., 17-/14-HDHA and 15-/12-HETE) in a distinct ratio which is specific for the different PUFA.^[59]^ The ratios in the supplemented macrophages (n-6/n-9 ratio: DHA, 2.6 ± 0.3; EPA, 6.4 ± 0.8) were similar, but slightly higher than the ratios reported for the isolated enzyme (n-6/n-9 ratio: DHA, 1.5; EPA 5.4).^[59]^ In line with this, chiral analysis of 15-HETE and 17-HDHA revealed that the (*S*)-product is predominantly present in the cells whereas the medium contains the racemate **(Figure S7)** supporting that the majority of the 15-LOX products in the cells were formed by enzymatic conversion from PUFA.

Considering other monohydroxy PUFA such as 4-HDHA, which can be formed by 5-LOX activity, concentration differences were comparable to oxylipins which can be only formed by autoxidation such as 10- or 13-HDHA **(Figure 3B top)**. Similar results were observed for EPA-derived 5-HEPE **(Figure S6)** and ARA-derived 5-HETE **(Figure 3B bottom)**. In contrast to the medium, cells contained mainly the (*S*)-enantiomers of 5- and 8-HETE **(Figure S7)**. This could indicate a selective uptake of the (*S*)-enantiomers from the medium or a formation of 5(*S*)-HETE by 5-LOX activity in the M2-like macrophages. However, only poor 5-LOX abundance and activity was reported for M2-like macrophages.^[29]^

The concentration of 10(*S*),17(*S*)-DiHDHA (*t*,*c*,*t*) (protectin (P)Dx) was approximately tenfold higher in the supplemented cells than in the non-supplemented cells **(Figure 3B top)**, whereas concentrations in the media were comparable **(Figure 3A top)**. For double hydroxylated products from ARA such as 8(*S*),15(*S*)-DiHETE and 5(*S*),15(*S*)-DiHETE, concentrations were higher in the medium with cells compared to medium without cells, but decreased in the cells following supplementation. This indicates that a formation of double hydroxylated oxylipins in the cells is likely caused by 15-LOX activity converting mono-hydroxylated oxylipins formed by autoxidation, as the levels of the dihydroxy-FA follow the changes of the monohydroxylated precursor. This was supported by chiral analysis **(Figure S7)**: 10-HDHA was present as a racemate in the medium and in the high supplemented macrophages indicating a formation by autoxidation and/or uptake from the medium. For 10,17-DiHDHA multiple isomers were detected including the 10(*S*),17(*S*)-DiHDHA (*t*,*c*,*t*) (PDx) and 10(*R*),17(*S*)-DiHDHA (*t*,*c*,*t*), indicating their formation by the conversion of racemic 10-HDHA by 15-LOX activity. 10(*R*),17(*S*)-DiHDHA (*t*,*t*,*c*) (neuroprotectin (NP)D1) coelutes with other isomers in reversed-phase chromatography **(Figure S7)**, as previously described.^[61]^ Thus, it is difficult or even impossible to support a relevant formation of NPD1 in the cells. For 5,15-DiHETE and 7,17-DiHDHA only one isomer, corresponding to the (*S*),(*S*)-product, was detected by chiral chromatography in the cells.^[43]^ Thus, the formation of double hydroxylated DHA and ARA oxylipins in M2-like macrophages is dependent on 15-LOX-1 activity as well as the amount of added mono-hydroxylated precursors/PUFA. Other dihydroxy- or trihydroxy-DHA belonging to the class of so-called “specialized pro-resolving mediators” (7-*epi*-maresin (MaR)1, MaR1, MaR2, RvD1, RvD2, RvD3), were not detected in the cells. Retention times and/or the ratio of different transitions did not match those of the corresponding authentic reference standard **(Figure S8)**. This is consistent with previous reports.^[62]^

Concentrations of 19(20)-EpDPE were higher in the medium without cells (high: 25.2 ± 2.6 nM) compared to medium with cells (high: 4.9 ± 0.5 nM) **(Figure 3A top)**. Consistently, the concentrations were strongly and dose-dependently increased in the supplemented macrophages **(Figure 3B top)** suggesting an uptake from the medium and incorporation into the cells. Here, for example, epoxy-PUFA are bound in phospholipids, as oxylipins were mainly found in the esterified form **(Figure 2D)**. In contrast, 19,20-DiHDPE concentrations were higher in the cells’ supernatant than in the medium without cells **(Figure 3B top)**. This could be explained by the conversion of 19(20)-EpDPE by soluble epoxide hydrolase activity within the cells and subsequent release into the medium. 19,20-DiHDPE was not detected in the cells. Following supplementation, increased concentrations of DHA-derived epoxy-FA and *vic*-dihydroxy-FA were described in human plasma^[56]^ whereby epoxy-PUFA were mainly esterified and dihydroxy-PUFA non-esterified.^[57]^

Overall, the results indicate that the observed shift in the oxylipin pattern following supplementation is either caused by oxylipin formation within the cells (e.g., 15-LOX products) or uptake of oxylipins from the media in a dose-dependent manner (e.g., 19(20)-EpDPE). Absolute levels of 15-LOX-1 oxylipins such as 17-HDHA or 15-HETE were changed the most by supplementation compared to other oxylipins. This indicates that 15-LOX-1 activity is mainly responsible for the formation of these oxylipins in supplemented M2-like macrophages. Here, the increased DHA- and EPA-derived oxylipins were at the expense of ARA-derived ones. This shift in the oxylipin pattern is in line with that reported in supplementation studies in mice and humans supporting that the *ex vivo* n-3 PUFA supplementation strategy allows the investigation of the effects of the n-3 PUFA in immune cells.

The change in the PUFA profile leads to a shift of oxylipins towards n-3 PUFA products which may contribute to the beneficial effects of n-3 PUFA on human health. Many oxylipins derived from n-3 PUFA such as 17-HDHA and 10,17-HDHA, which are massively increased by supplementation, are regarded as anti-inflammatory lipid mediators.^[63–66]^ In contrast, oxylipins from ARA, which decrease following supplementation, are predominantly described having pro-inflammatory effects such as fever, pain or bronchoconstriction.^[17]^ However, further research is needed to investigate the role of oxylipins in the physiological effects of n-3 PUFA supplementation. The developed *ex vivo* supplementation strategy is a promising tool for these experiments.

### Effects of n-3 PUFA supplementation on LPS stimulation of macrophages

The developed *ex vivo* supplementation strategy was used for investigating changes in the response of the macrophages to the inflammatory stimulus LPS. The LPS stimulus resulted in a highly elevated induction (100-700 fold) of the mRNA levels of pro-inflammatory cytokines such as tumor necrosis factor (TNF)α or IL-6 **(Figure S9)**. However, n-3 PUFA supplementation did not significantly change the expression of cytokines and chemokines, with or without LPS stimulation **(Figure 4A-B)**. This is in contrast to *in vitro* experiments, which reported a downregulation of inflammatory genes following n-3 PUFA supplementation.^[67–69]^ However, in human supplementation studies, the results regarding inflammatory cytokines are inconsistent,^[23]^ highlighting the need for further studies in order to investigate the mechanism of how n-3 PUFA impact inflammatory processes.

**Figure 4:**
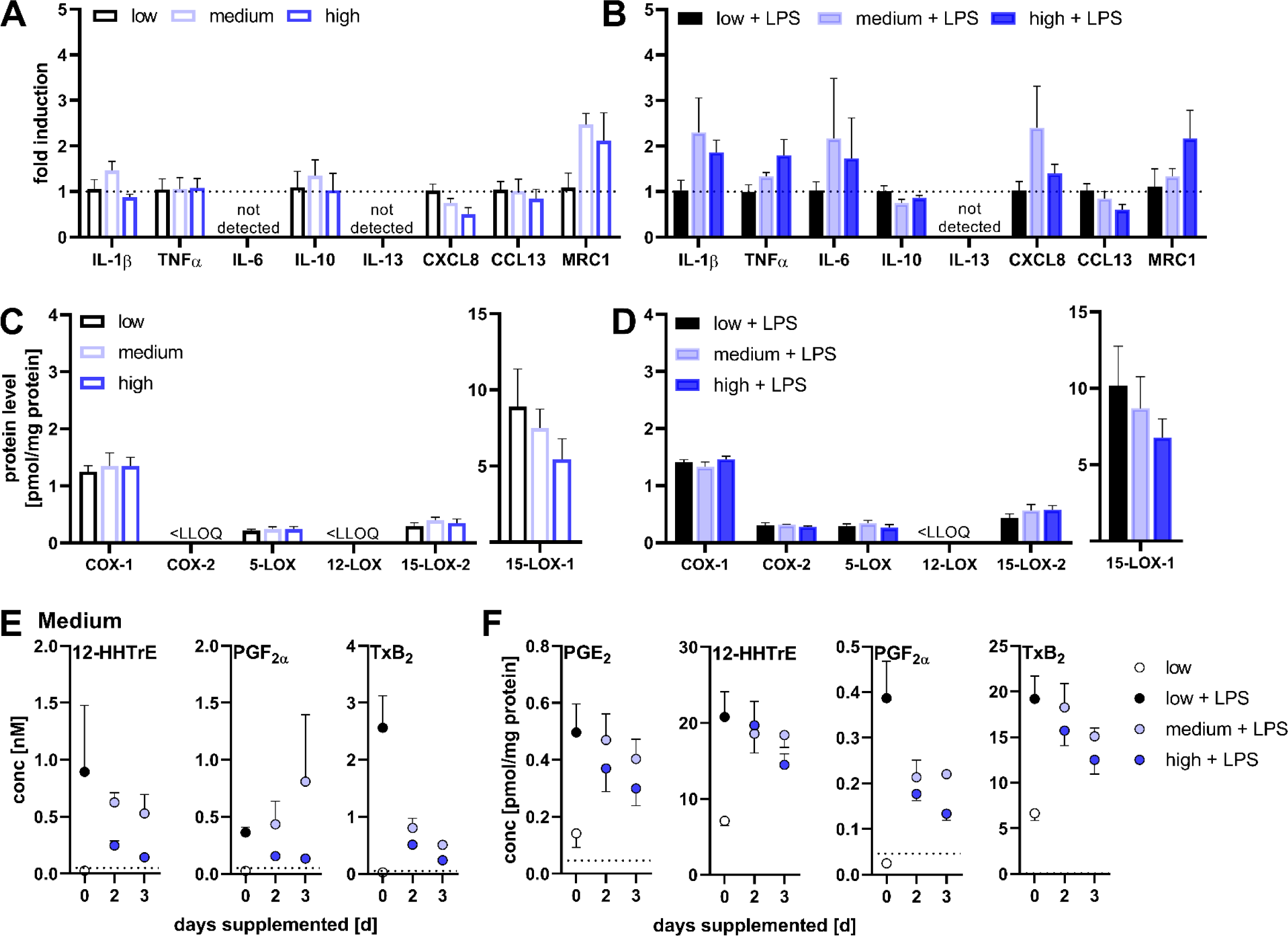
LPS elicits changes in supplemented vs. non-supplemented M2-like macrophages. After isolation of monocytes and differentiation into M2-like macrophages, cells were supplemented using media containing 4 µM (low, non-supplemented control), 21 µM (medium) or 41 µM (high) DHA. For LPS stimulation 1 µg mL^-1^ LPS was added during the final 6 h. Following 3 days of supplementation, changes in **A-B)** the transcription of selected genes analyzed by qPCR and **C-D)** abundance of ARA cascade enzymes determined by quantitative proteomics. Non-esterified oxylipins in **E)** Medium and **F)** cell pellets determined by LC-MS/MS. Results are shown as mean ± SEM or SD, *n* = 3. Dotted line indicates the lower limit of quantification (LLOQ).

The macrophage marker mannose receptor C-type 1 (MRC1) tended to increase by n-3 PUFA supplementation **(Figure 4A-B)**, whereas protein levels of IL1R2, toll-like receptor 2 and toll-like receptor 4 were not changed, indicating only minor or no changes of proteins involved in the polarization of macrophages by n-3 PUFA or LPS. The levels of ARA cascade enzymes such as COX-1, 5- and 15-LOX-2 were low (< 2 pmol mg^-1^ protein), whereas 15-LOX-1 was highly abundant (8.9 ± 2.5 pmol mg^-1^ protein) in the M2-like macrophages **(Figure 4C)**. This is in line with previously described levels.^[29, 31]^ LPS did not affect the levels of these enzymes **(Figure 4D)**. Hence, oxylipins formed by these pathways or autoxidation were less affected by LPS stimulation and the oxylipin pattern was comparable to that of unstimulated macrophages **(Figure S10)**.

LPS stimulus resulted in massive elevated COX-2 levels in the macrophages **(Figure 4D)**. In line with this, concentrations of COX products such as prostaglandins were increased in medium and cells **(Figure 4E-F, S10)**. Similarly, concentrations of 11- and 15-HETE, also products of COX-2, were increased after LPS stimulation. These were reduced by DHA supplementation in a dose- and time-dependent manner **(Figure S10)**. Pro-inflammatory prostaglandins such as PGE_2_ and TxB_2_ were also reduced in the macrophages and medium after DHA supplementation, e.g. PGE_2_ by about 40% in the cells and TxB_2_ by about 90% in the medium **(Figure 4E-F)**. Similarly, supplementation studies reported decreased levels of prostaglandins following n-3 PUFA supplementation.^[70–73]^ Our finds support a reduced conversion of ARA by COX-2 following n-3 PUFA supplementation, which results in lower concentrations of pro-inflammatory prostaglandins which is one of the major mechanisms how the anti-inflammatory effect of n-3 PUFA is mediated.

## Concluding remarks

Overall, our experiments showed how primary human macrophages can be supplemented with DHA to alter the FA and oxylipin profile, mimicking n-3 PUFA supplementation studies in humans. As n-3 PUFA supplementation in cell culture experiments provides many challenges, key parameters were identified and controlled ensuring a reliable strategy. Autoxidation of PUFA cannot be completely avoided, but the extent can be reduced and should be carefully monitored during the process. This is particularly important because several oxidized PUFA are biologically active lipid mediators, and thus, low concentration might interfere with investigated inflammatory processes. Here, we present an *ex vivo* supplementation strategy for primary human macrophages addressing these challenges. The observed changes in the PUFA profile and the shift in the oxylipin pattern towards n-3 PUFA are in line with intervention studies in humans. Applying the model to investigate the inflammatory stimulus LPS, decreased levels of pro-inflammatory prostaglandins were observed in the DHA-supplemented macrophages which may be one of the mechanisms by which n-3 PUFA mediate their anti-inflammatory effects. Thus, the developed *ex vivo* supplementation strategy is a helpful tool for mechanistic investigations of n-3 PUFA and oxylipins in primary human immune cells.

## Conclusion

The developed *ex vivo* n-3 PUFA supplementation strategy allows to change the FA pattern of monocytes from subjects with a typical Western Diet to macrophages with a FA pattern comparable to subjects having a high n-3 PUFA status. By controlling key parameters for selection of buffy coats, plasma and n-3 PUFA preparation as well as preparation of the medium, a reliable supplementation is possible, allowing to alter the FA and oxylipin patterns of macrophages comparable to human intervention studies. Thus, this tool allows mechanistic investigation of long-chain n-3 PUFA and arising oxylipins in primary human immune cells under strictly controlled conditions without the need for human intervention studies.

## Declarations

### Ethics approval

Peripheral blood monocytic cells (PBMC) were isolated from buffy coats obtained from blood donations. Blood samples were drawn with the informed consent of the human subjects. The study was approved by the Ethical Committee of the University of Wuppertal.

## Abbreviations

ARA: arachidonic acid
COX: cyclooxygenase
CXCL8: C-X-C motif ligand 8
DHA: docosahexaenoic acid
DiHDHA: dihydroxy docosahexaenoic acid
DiHETE: dihydroxy eicosapentaenoic acid
EPA: eicosapentaenoic acid
EpDPE: expoxy docosapentaenoic acid
FA: fatty acid
GM-CSF: granulocyte-macrophage colony-stimulating factor
HDHA: hydroxy docosahexaenoic acid
HEPE: hydroxy eicosapentaenoic acid
HETE: hydroxy eicosatetraenoic acid
HUFA: highly unsaturated fatty acid
IS: internal standard
LOX: lipoxygenase
MaR: maresin
M-CSF: macrophage colony-stimulating factor
MRC1: mannose receptor C-type 1
MUFA: monounsaturated fatty acid
NP: neuroprotectin
PBMC: peripheral blood monocytic cells
P: protectin
PG: prostaglandin
PUFA: polyunsaturated fatty acid
P/S: penicillin/streptomycin
Rv: resolvin
SFA: saturated fatty acid
THP-1: human monocyte cell line
TNFα: tumor necrosis factor α
Tx: thromboxane

## Supporting Information

**Table S1:**
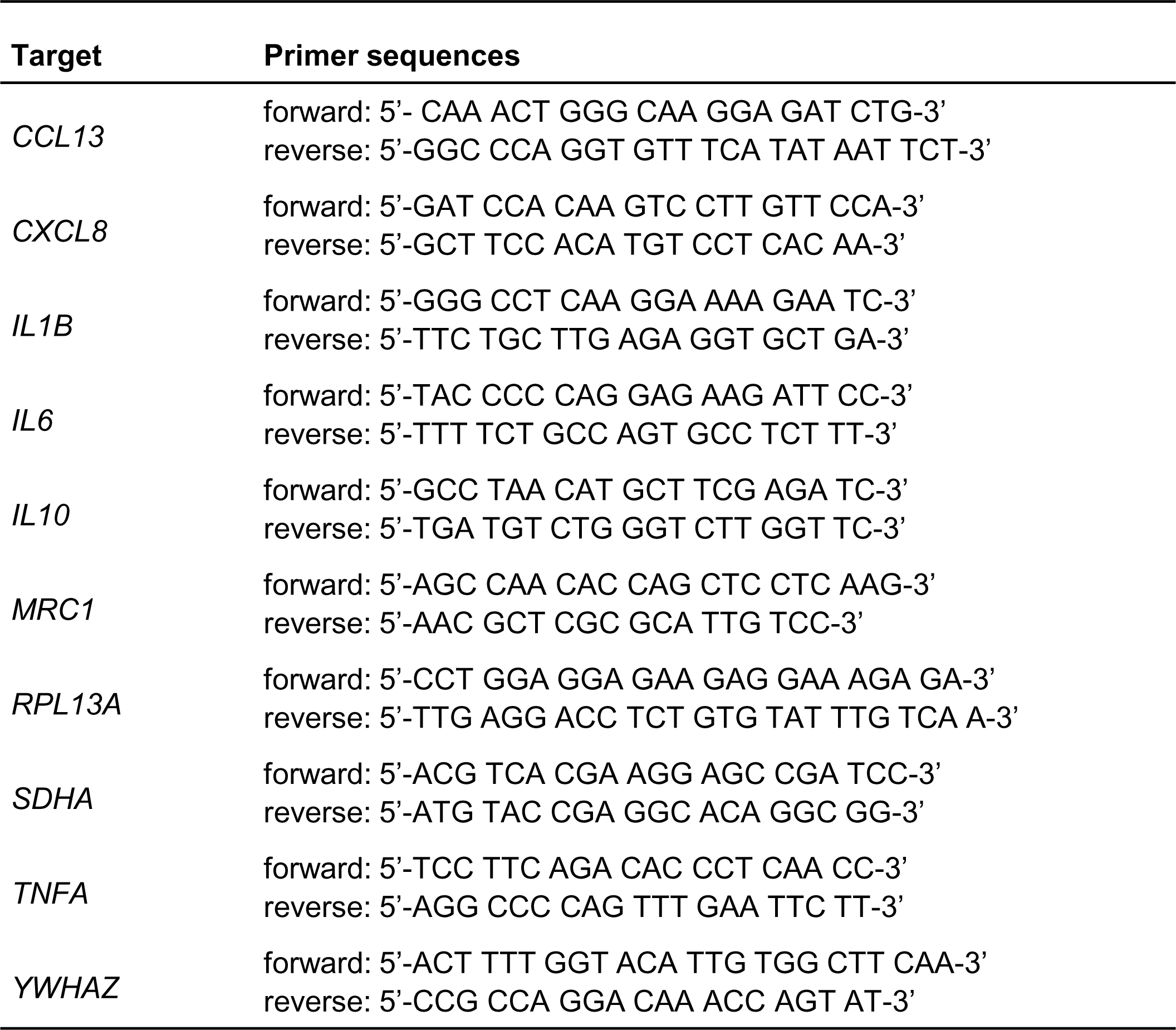
Primer sequences used for quantitative real-time PCR.

**Figure S1:**
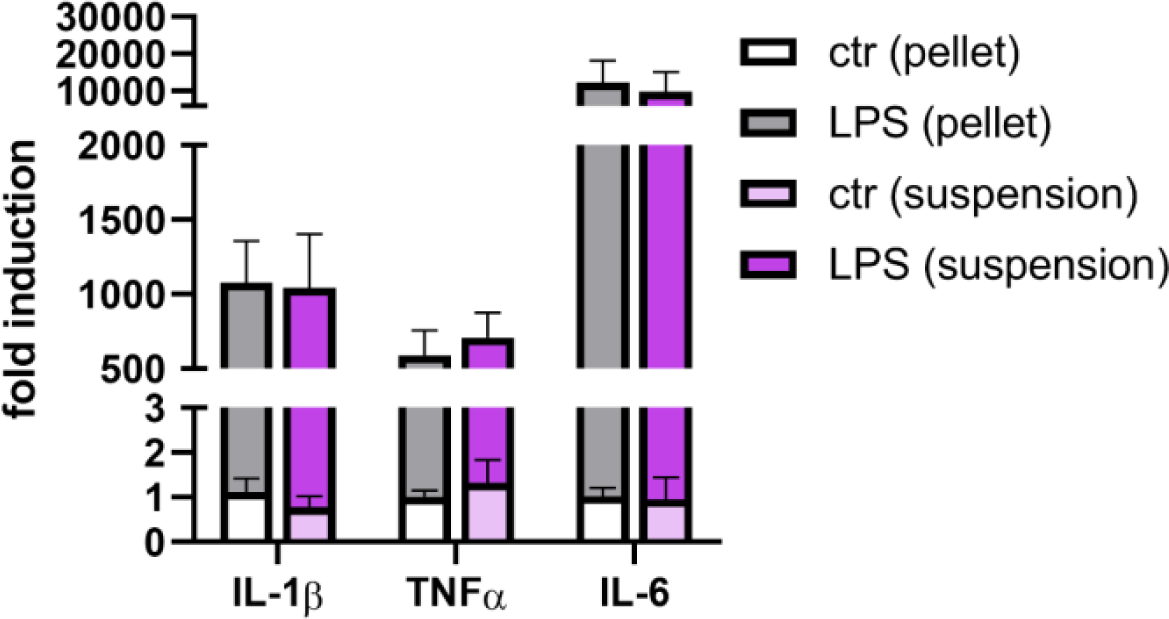
Relative mRNA expression of macrophages using different sample preparation methods. Primary human monocytic cells were differentiated into macrophages using 10 ng mL^-1^ M-CSF for 7 days and additional 10 ng mL^-1^ IL-4 for the final 48 h. For LPS stimulation 1 µg mL^-1^ LPS was added 6 h before harvest. Cells were harvested by cold shock method and either stored as dry pellet (pellet) or resuspended in PBS, homogenized by sonication and stored as cell suspension in PBS (suspension) at −80 °C followed by mRNA analysis. Results are shown as mean ± SEM, *n* = 3 subjects.

**Figure S2:**
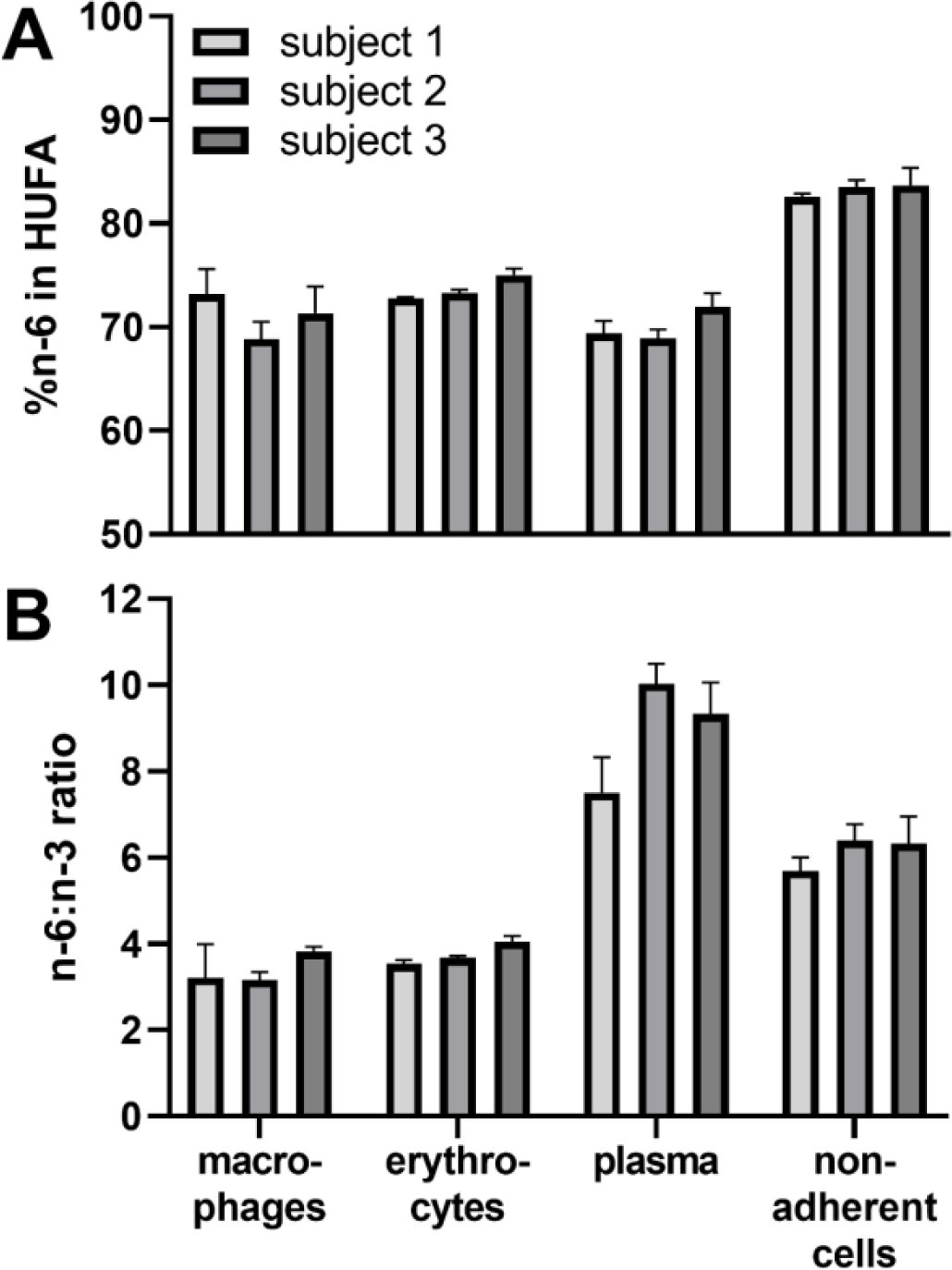
Characterization of FA status of three exemplarily human subjects based on A) %n-6 in HUFA and B) n-6:n-3 ratio of buffy coat fractions. Fractions from buffy coats of three subjects were collected: erythrocytes after dextran sedimentation, plasma after density gradient centrifugation and remaining non-adherent cells after monocytic cells getting adherent for 1-2 h. Monocytic cells were differentiated with 10 ng mL^-1^ M-CSF for 7 days and additional 10 ng mL^-1^ IL-4 for final 48 h into M2-like macrophages. Cell pellets and plasma were analyzed for total fatty acid concentrations by LC-MS/MS. A) %N-6 in HUFA was calculated from total fatty acid concentrations of C20:3 n-6, C20:4 n-6, C22:4 n-6, C22:5 n-6, C20:3 n-9, C20:5 n-3, C22:5 n-3, C22:6 n-3 by LC-MS and B) n-6:n-3 ratio was calculated from C18:3 n-3, C18:2 n-6, C20:4 n-6, C20:5 n-3 and C22:6 n-3. Results are shown as mean ± SD, *n* = 3.

**Table S2:** Fatty acid status of the human subjects. Erythrocytes were analyzed for total fatty acid concentrations by means of LC-MS/MS and %n-6 in HUFA was calculated from fatty acid concentrations of C20:3 n-6, C20:4 n-6, C22:4 n-6, C22:5 n-6, C20:3 n-9, C20:5 n-3, C22:5 n-3, C22:6 n-3. All 12 donors were healthy, voluntary blood donors from a local blood donation center. Results are shown as mean ± SD, *n* = 3.

| Subject | %n-6 in HUFA |
| --- | --- |
|  | [%] |
| 1 | 72.8 ± 0.2 |
| 2 | 73.3 ± 0.3 |
| 3 | 75.0 ± 0.2 |
| 4 | 75.1 ± 0.3 |
| 5 | 74.7 ± 0.4 |
| 6 | 72.2 ± 1.0 |
| 7 | 76.7 ± 0.2 |
| 8 | 75.1 ± 1.3 |
| 9 | 75.1 ± 1.0 |
| 10 | 82.9 ± 0.6 |
| 11 | 77.7 ± 0.3 |
| 12 | 74.6 ± 0.6 |

**Table S3:**
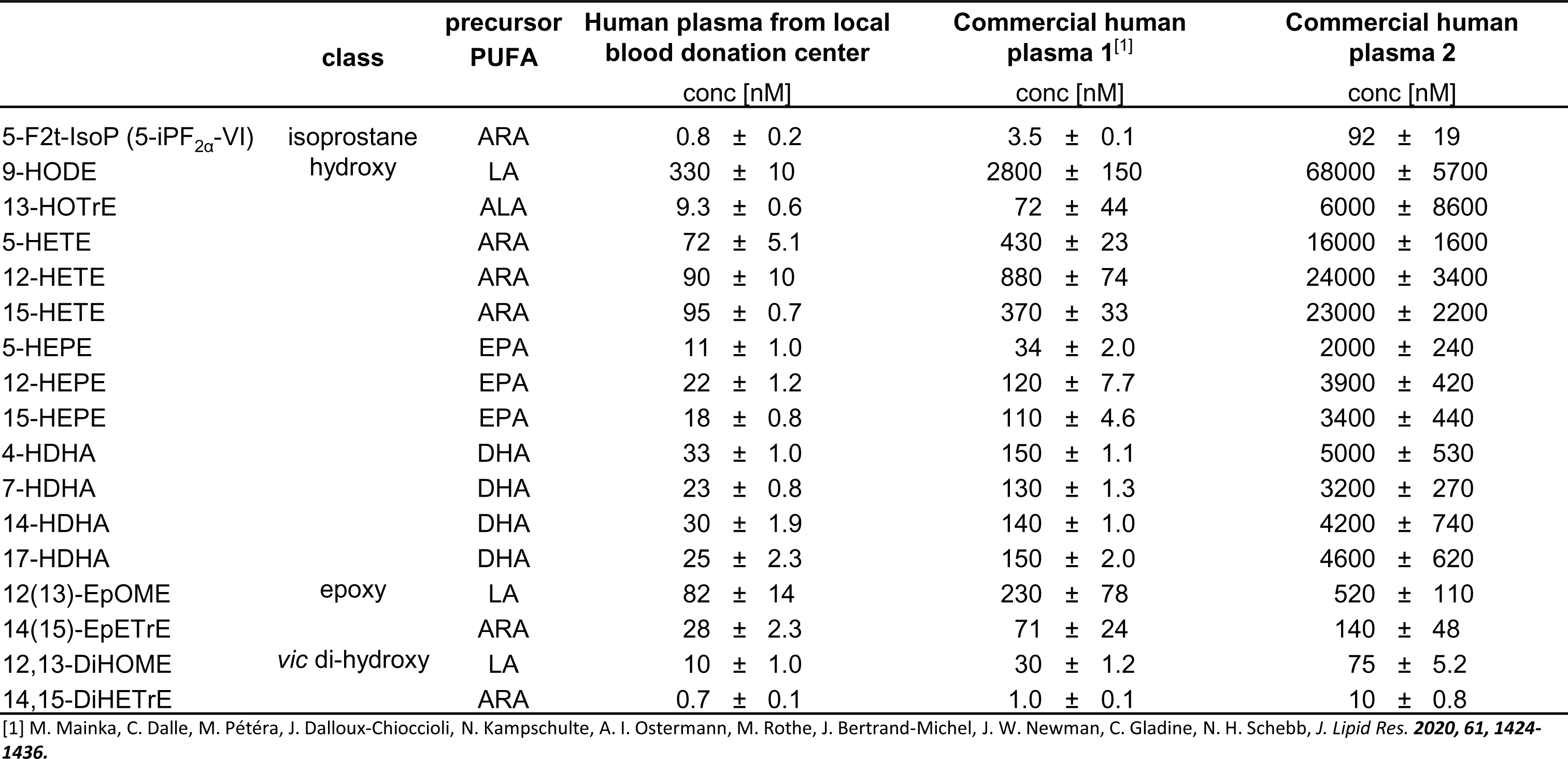
Oxylipin levels in different human plasmas. Human plasma from a local blood donation center (pool of 4 subjects) as well as two commercially available human plasmas were analyzed for total oxylipin concentrations by means of LC-MS/MS. Representative oxylipins derived from different precursor PUFA covering different classes are shown as mean ± SD, *n* = 3.

**Figure S3:**
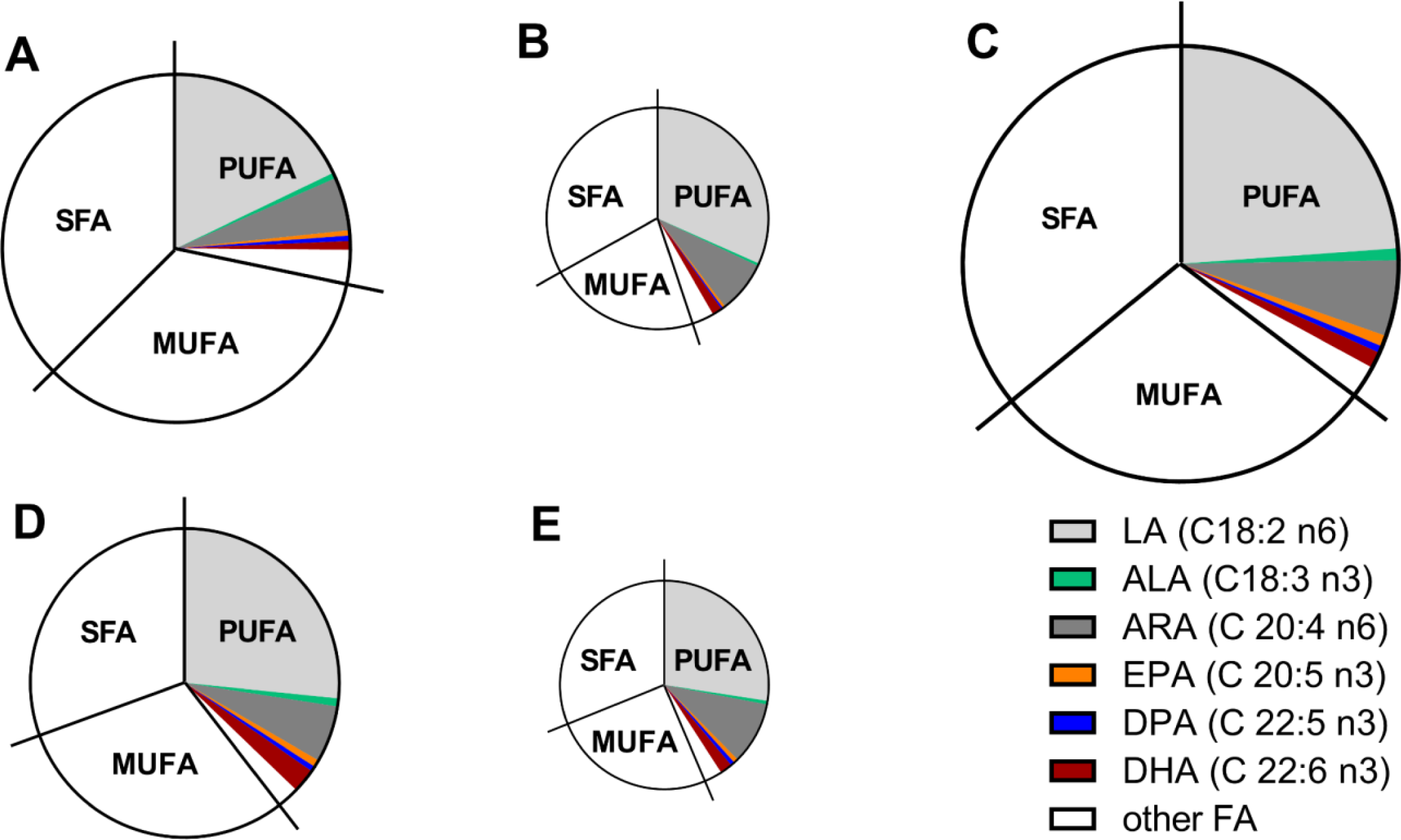
Relative fatty acid profiles of human plasmas. Non-fasting plasma of five different subjects from a local blood donation center was analyzed for total FA concentrations by means of GC-FID. Sizes of the circles indicate total fatty acid concentrations relatively on plasma’s C total fatty acid concentration (plasma A, 11.9 ± 1.3 mM, 80%; plasma B, 7.7 ± 0.1 mM, 52%; plasma C, 14.9 ± 1.1 mM, 100%; plasma D, 10.6 ± 0.3 mM, 71%; plasma E, 7.1 ± 0.4 mM, 48%). SFA, saturated fatty acids; MUFA, monounsaturated fatty acids; PUFA, polyunsaturated fatty acids

**Table S4:**
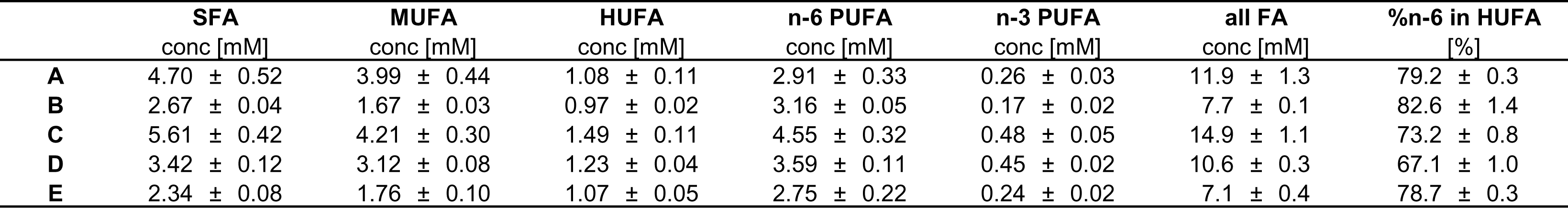
Fatty acid pattern of human plasmas. Non-fasting plasma of five different subjects from a local blood donation center was analyzed by means of GC-FID. Results are shown as mean ± SD, *n* = 3. %N-6 in HUFA was calculated from fatty acid concentrations of C20:3 n-6, C20:4 n-6, C22:4 n-6, C22:5 n-6, C20:3 n-9, C20:5 n-3, C22:5 n-3, C22:6 n-3. SFA, saturated fatty acids; MUFA, monounsaturated fatty acids; HUFA, highly unsaturated fatty acids; PUFA, polyunsaturated fatty acids

**Table S5:** Purity of commercial fatty acid preparations. Different free fatty acid preparations were analyzed for oxylipin concentrations by means of LC-MS/MS. Shown are mean ± SD, *n* = 3.

|  | **Oxylipins** |
| --- | --- |
|  | **[% (w/w)]** |
| DHA > 99% | 0.36 ± 0.01 |
| EPA > 99% | 0.21 ± 0.02 |
| EPA ≥ 97% | 3.30 ± 0.14 |

**Table S6:**
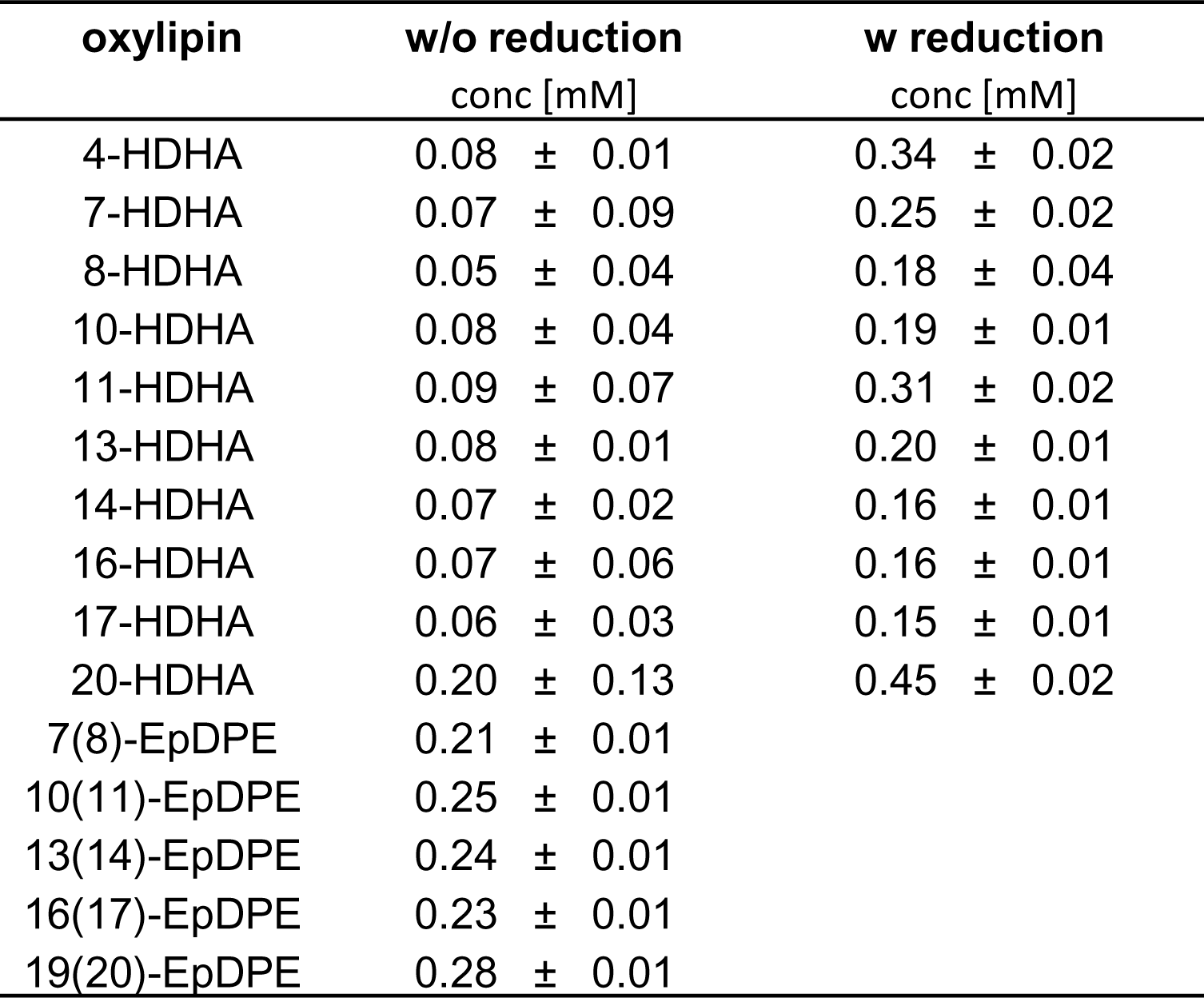
Oxylipins in commercial n-3 PUFA preparation used for supplementation without (w/o) and with (w) reduction. DHA >99% (1 M) was analyzed for oxylipin concentrations by means of LC-MS/MS. Reduction of hydroperoxyl-FA to hydroxy FA was carried out using SnCl_2_ (10 mg mL^-1^). Shown are mean ± SD, *n* = 3.

| oxylipin | w/o reduction | w reduction |
| --- | --- | --- |
|  | conc [mM] | conc [mM] |
| 4-HDHA | 0.08 ± 0.01 | 0.34 ± 0.02 |
| 7-HDHA | 0.07 ± 0.09 | 0.25 ± 0.02 |
| 8-HDHA | 0.05 ± 0.04 | 0.18 ± 0.04 |
| 10-HDHA | 0.08 ± 0.04 | 0.19 ± 0.01 |
| 11-HDHA | 0.09 ± 0.07 | 0.31 ± 0.02 |
| 13-HDHA | 0.08 ± 0.01 | 0.20 ± 0.01 |
| 14-HDHA | 0.07 ± 0.02 | 0.16 ± 0.01 |
| 16-HDHA | 0.07 ± 0.06 | 0.16 ± 0.01 |
| 17-HDHA | 0.06 ± 0.03 | 0.15 ± 0.01 |
| 20-HDHA | 0.20 ± 0.13 | 0.45 ± 0.02 |
| 7(8)-EpDPE | 0.21 ± 0.01 |  |
| 10(11)-EpDPE | 0.25 ± 0.01 |  |
| 13(14)-EpDPE | 0.24 ± 0.01 |  |
| 16(17)-EpDPE | 0.23 ± 0.01 |  |
| 19(20)-EpDPE | 0.28 ± 0.01 |  |

**Figure S4:**
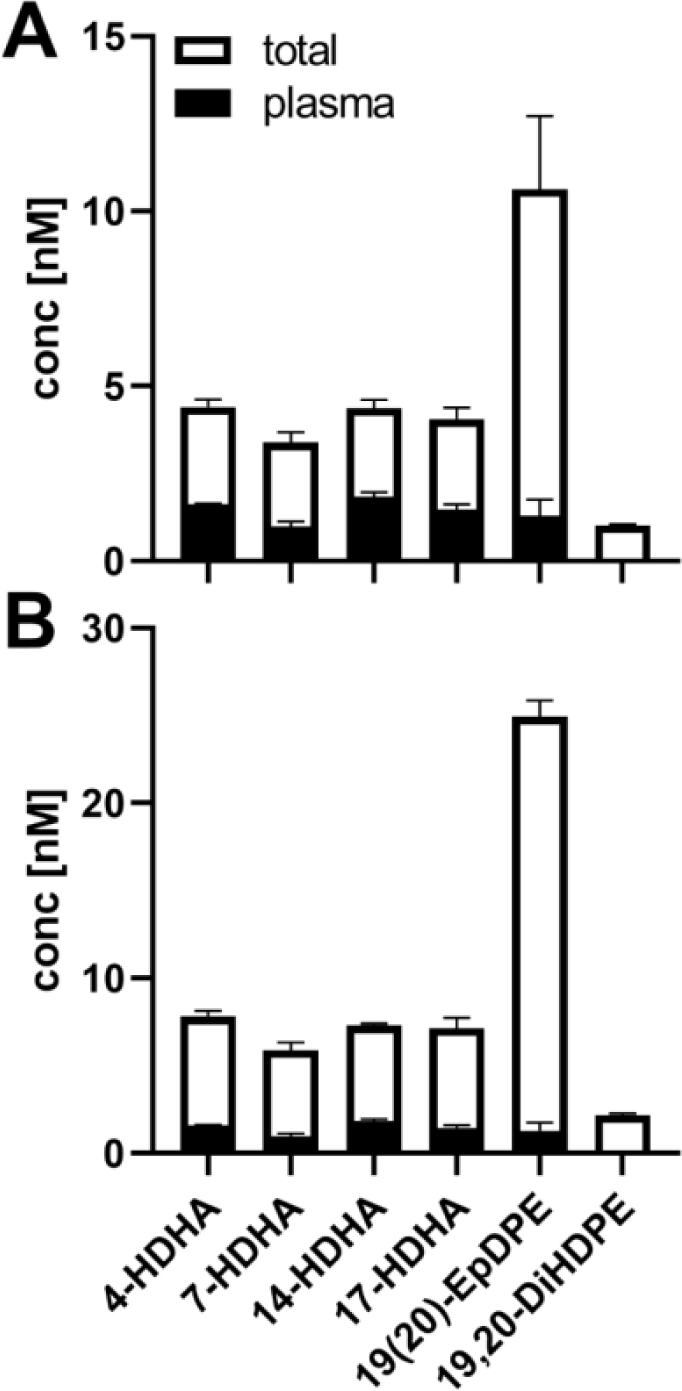
Human plasma DHA derived oxylipins in A) low and B) high supplemented medium. **A)** Low and **B)** high supplemented medium and plasma used were analyzed for total oxylipin concentrations. Plasma oxylipin concentrations were substracted proportionally from those of the medium. Results are shown as mean ± SD, *n* = 3.

**Figure S5:**
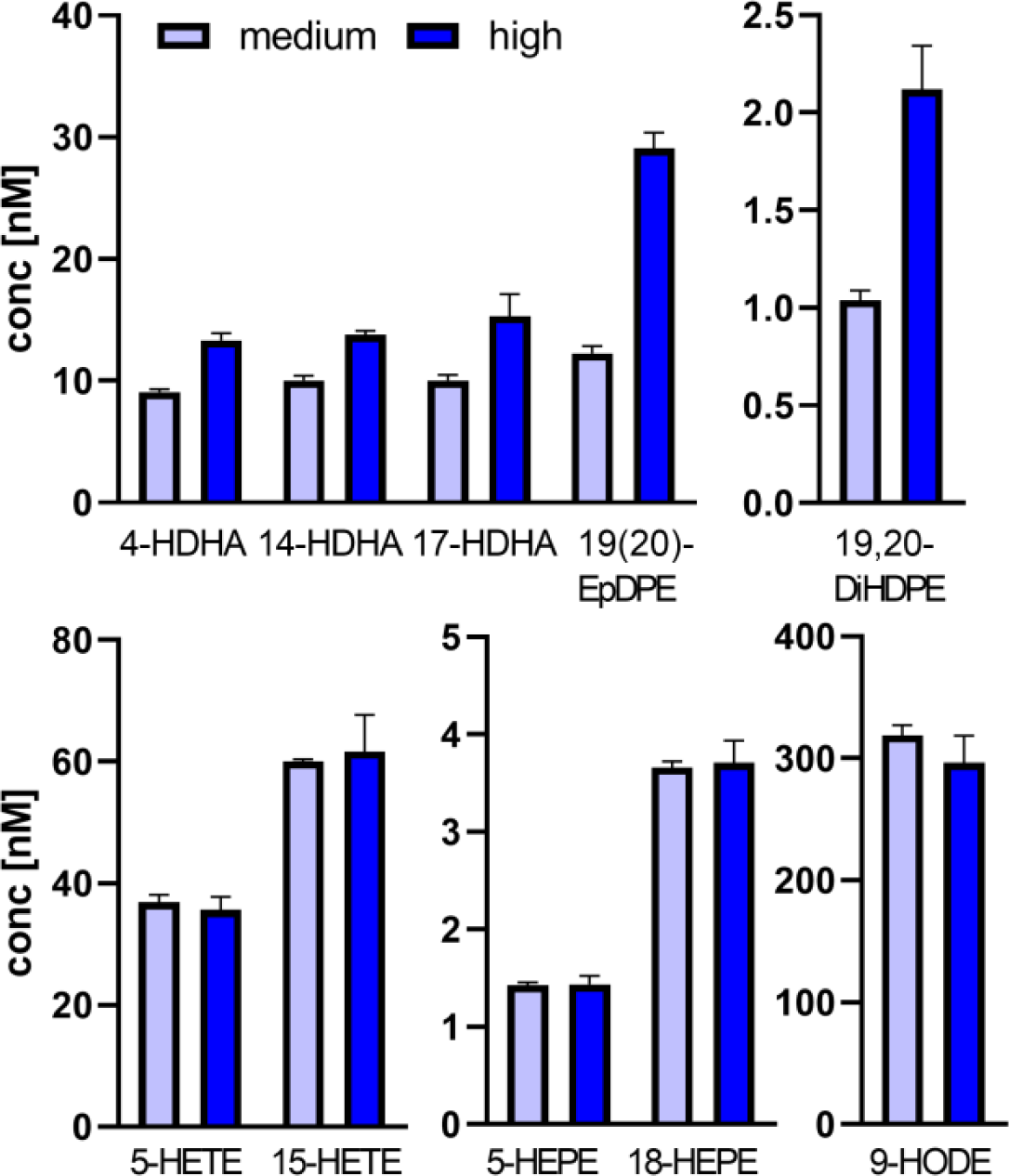
Oxylipin levels of supplemented media after storage at 4 °C for one day. Medium and high supplemented medium were stored in the fridge (4 °C) for one day and analyzed for total oxylipin concentrations by means LC-MS/MS. Shown are representative oxylipins derived from ARA, DHA and ALA. Results are shown as mean ± SD, *n* = 3.

**Figure S6:**
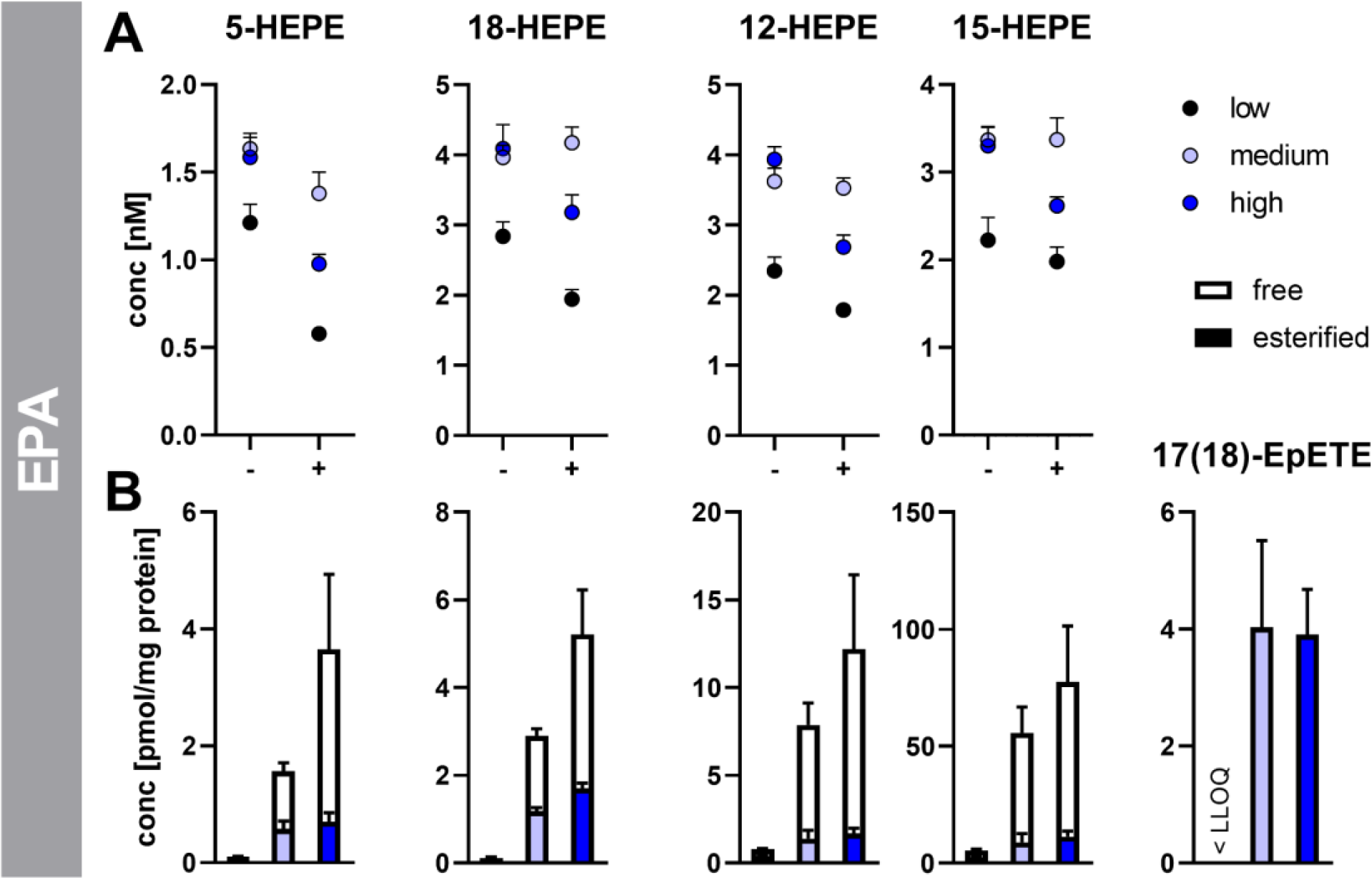
Oxylipin pattern of A) medium (without (-) and with cells (+)) and B) the macrophages after two days of incubation with different levels of DHA. Monocytes were isolated from buffy coats and differentiated into M2-like macrophages. Incubations were carried out using media containing 4 µM (low, non-supplemented control), 21 µM (medium) or 41 µM (high) DHA for 2 days with and without cells. Cell culture media was analyzed for total oxylipins, and cell pellets for total and non-esterified oxylipins by means of LC-MS/MS. Results are shown as mean ± SD (medium) or mean ± SEM (cells), *n* = 3.

**Figure S7:**
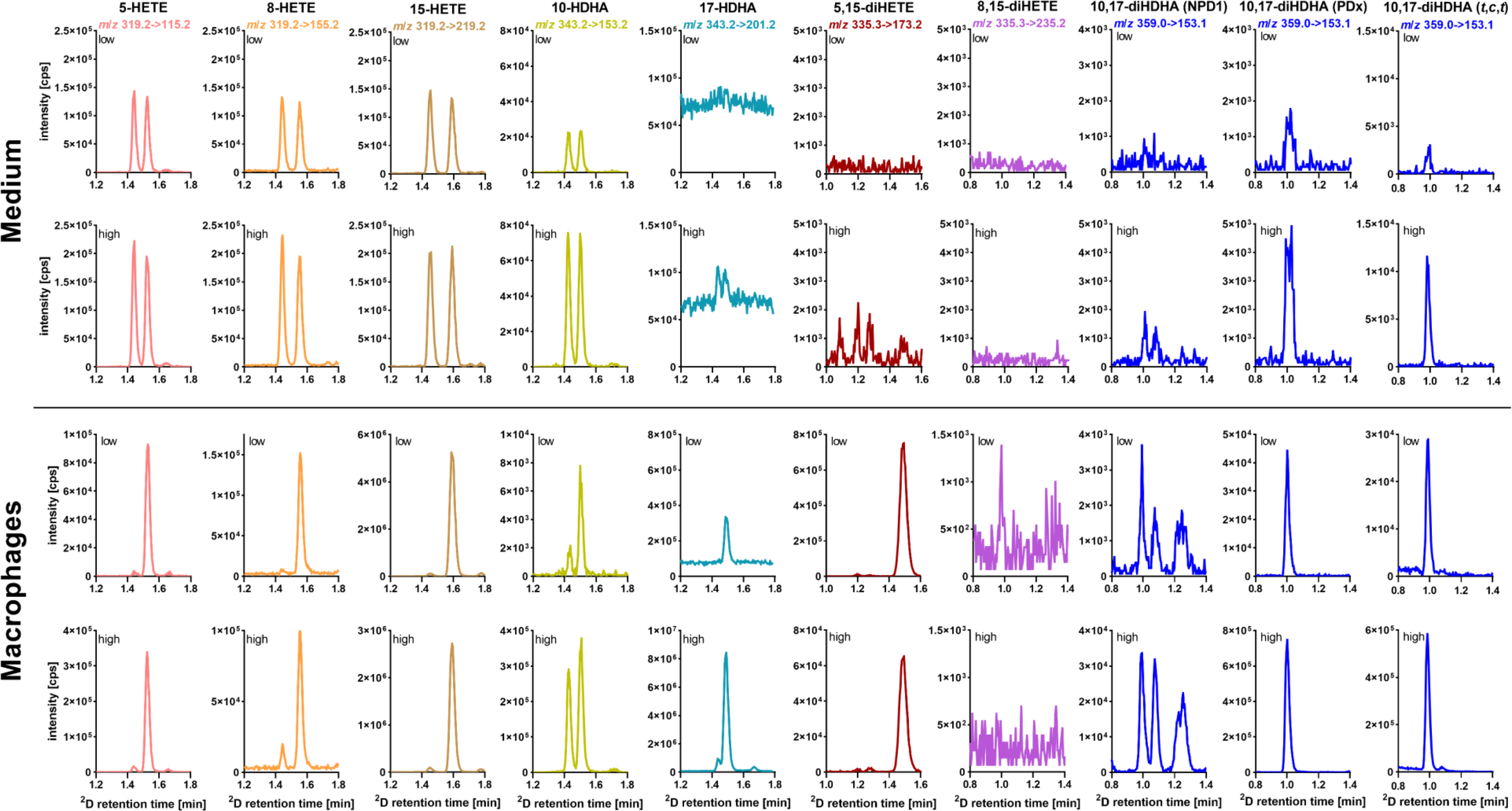
Chiral LC analysis of oxylipins in medium and macrophages. Media were prepared by adding no (low, non-supplemented control) or 45 µM (high) DHA preparation (< 99%) to cell culture medium. Primary human macrophages were supplemented using the different media for 2 days. Shown are the chromatograms of the chiral LC analysis of the indicated oxylipins in the cells and media stored at 37 °C for two days.

**Figure S8:**
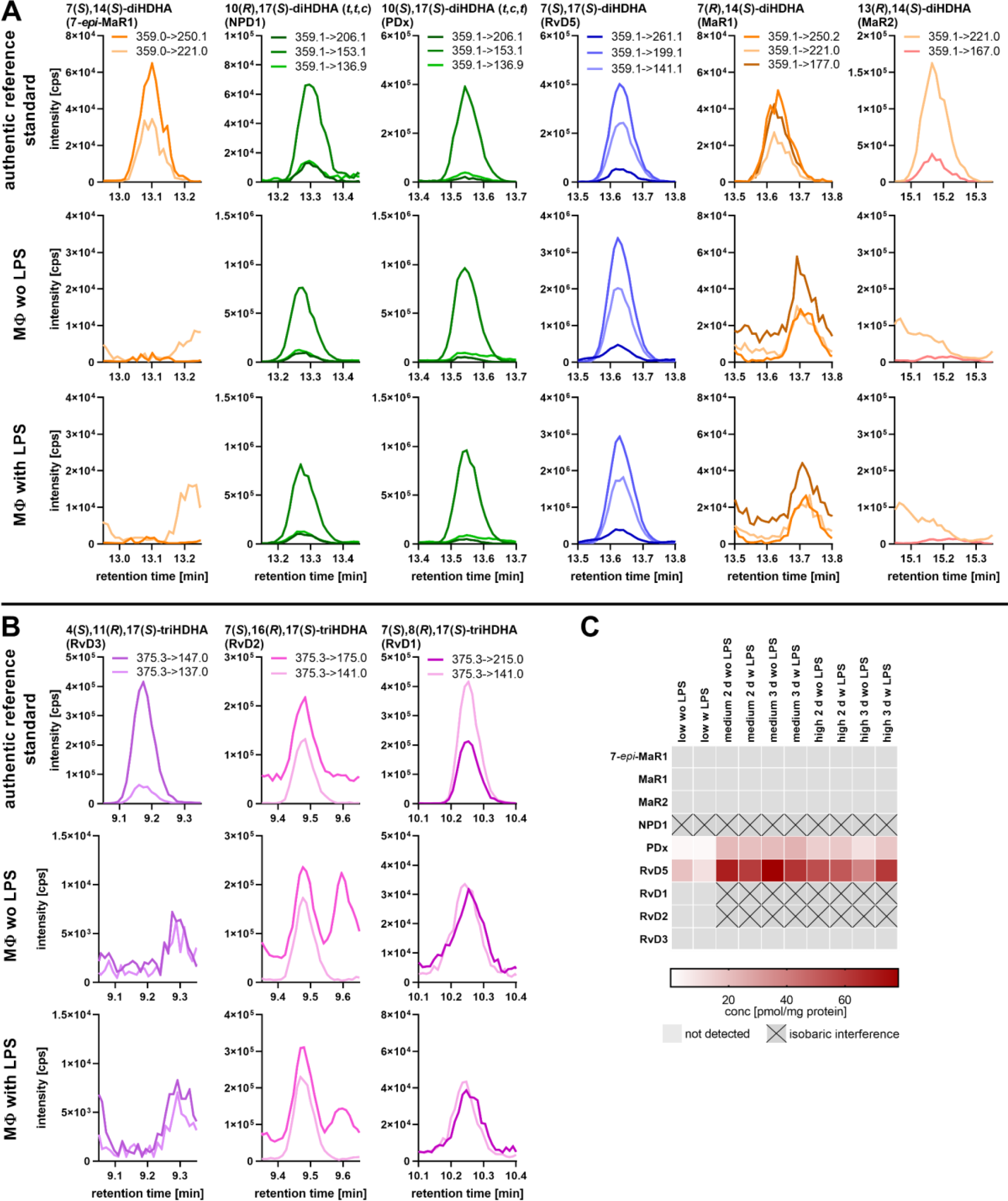
Analysis of di- and tri-OH-DHA oxylipins in DHA-supplemented macrophages. Primary human macrophages were supplemented using media containing 45 µM (high) DHA for 2 days without (wo) or with LPS stimulation (1 µg/mL) for 6 h. Shown are the MRM chromatograms (2-3 transitions) of **A)** di- and **B)** tri-OH-DHA oxylipins in the authentic reference standard and the macrophages (MΦ) without (wo) and with (w) LPS stimulation of one exemplarily subject and **C)** overview about analyzed di- and tri-OH-DHA in the macrophages.

**Figure S9:**
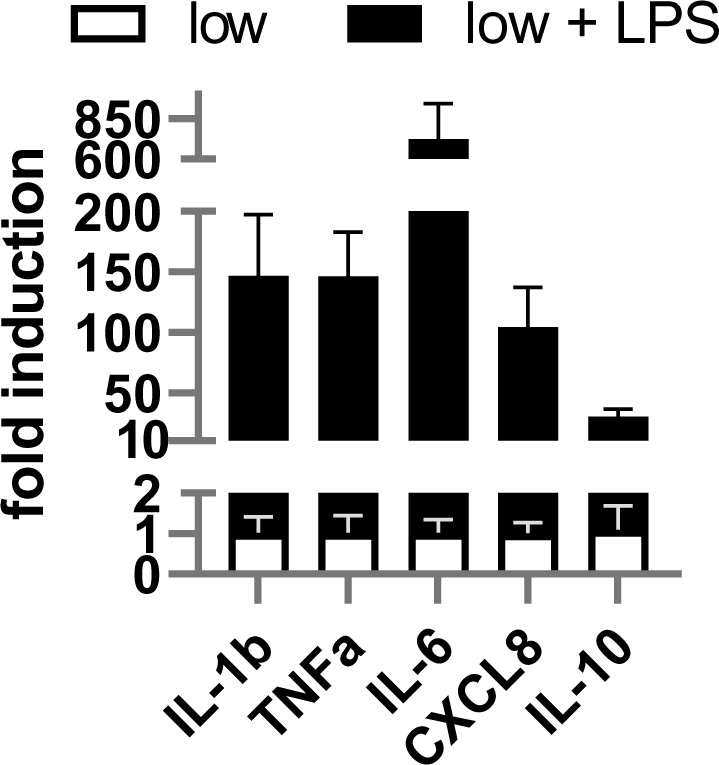
Relative mRNA induction of pro-inflammatory cytokines after LPS stimulus. Primary human monocytic cells were differentiated into macrophages using 10 ng mL^-1^ M-CSF for 7 days and additional 10 ng mL^-1^ IL-4 for the final 48 h. For LPS stimulation 1 µg mL^-1^ LPS was added into the medium 6 h before harvest. After harvest, cells were analyzed for mRNA levels by qPCR. Results are shown as mean ± SEM, *n* = 3 subjects.

**Figure S10:**
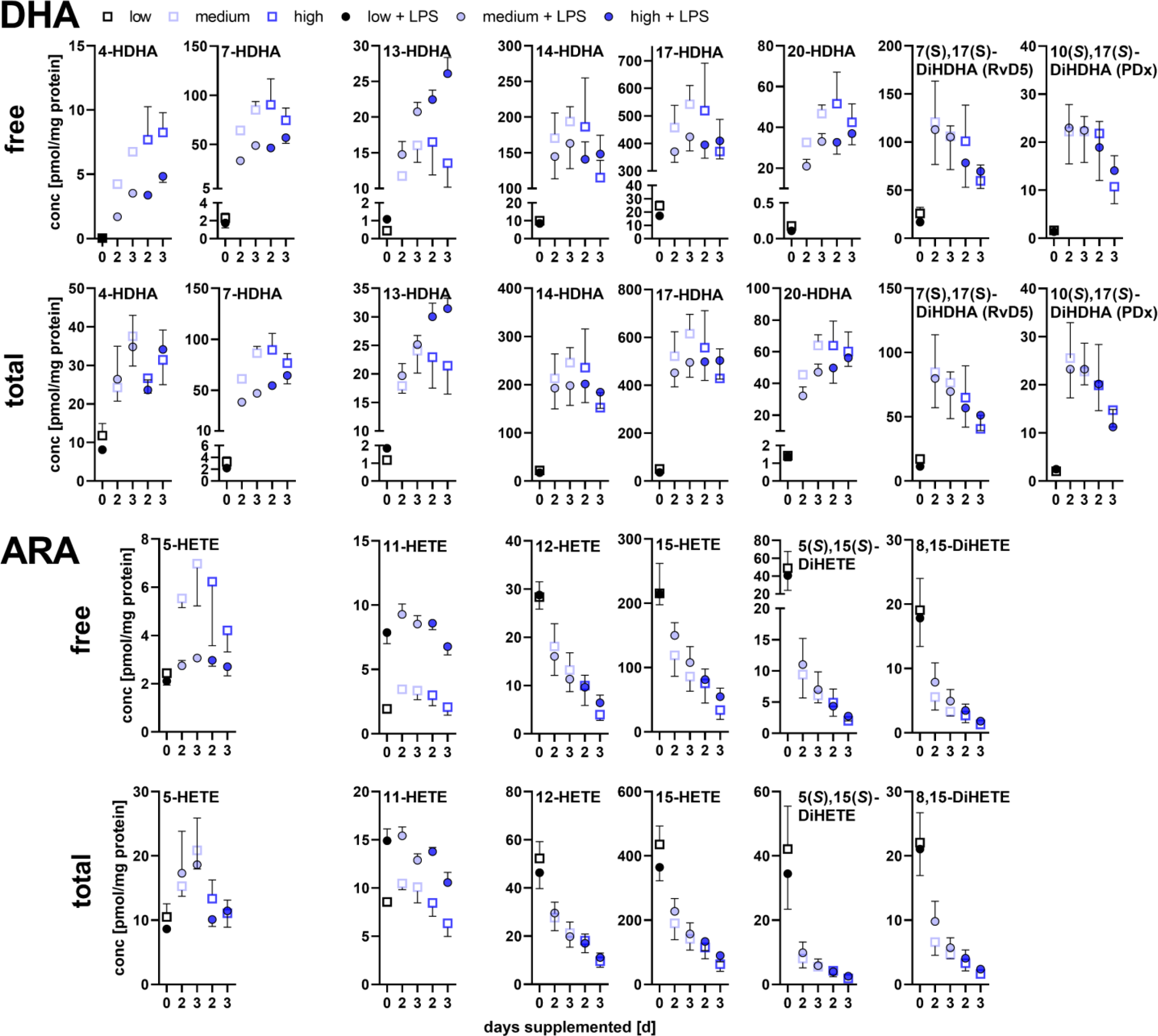
Free and total oxylipins in DHA-supplemented macrophages with and without LPS stimulus. Primary human monocytic cells were differentiated into macrophages using 10 ng mL^-1^ M-CSF for 7 days and additional 10 ng mL^-1^ IL-4 for the final 48 h. For LPS stimulation 1 µg mL^-1^ LPS was added into the medium 6 h before harvest. After harvest, cells were analyzed for total and free oxylipins by means of LC-MS/MS. Results are shown as mean ± SEM, *n* = 3 subjects.

